# Patient-Derived Synovium-on-a-Chip Recapitulates Disease and Treatment Response and Permits Study of Resistance in Rheumatoid Arthritis

**DOI:** 10.64898/2026.10.30.763140

**Authors:** Ruiqi Chen, Theresa Wampler Muskardin, Fanghao Shi, Yeji Lee, Christele Felix, Carolyn Harris, Lunan Liu, Eric Suh, Emily Wu, Scott Rodeo, David Mayman, Daniel Ramirez, Katsuo Kurabayashi, Timothy B. Niewold, Weiqiang Chen

**Affiliations:** Department of Biomedical Engineering, Tandon School of Engineering, New York University, Brooklyn, NY, USA; Department of Medicine, Hospital for Special Surgery, New York, NY, USA; Department of Medicine and Department of Pediatrics, Weill Cornell Medical College, New York, NY, USA; Department of Mechanical and Aerospace Engineering, Tandon School of Engineering, New York University, Brooklyn, NY, USA; Department of Orthopedic Surgery, Hospital for Special Surgery, New York, NY, USA; Department of Orthopedic Surgery, Weill Medical College of Cornell University, New York, NY, USA; Adult Reconstruction and Joint Replacement Service Department, Hospital for Special Surgery, New York, NY, USA; Pathology and Laboratory Medicine Department, Hospital for Special Surgery, New York, NY, USA; School of Biomedical Engineering, Nanjing University, Suzhou, China

## Abstract

Rheumatoid arthritis (RA) is a chronic inflammatory disease with substantial interpatient heterogeneity in synovial pathology and therapeutic response. Reliable methods to predict drug efficacy for a given patients remain a critical unmet need. We developed a vascularized, immunocompetent synovium-on-a-chip platform that integrates patient-derived fibroblast-like synoviocytes (FLS) with matched peripheral blood leukocytes to provide a physiological human model of RA. The platform reconstructed the lining–sublining synovial architecture, recapitulated key features of RA inflammation, and exhibited transcriptional profiles resembling those of patient synovial tissue. Across patients spanning low, mixed, and high inflammatory endotypes, stromal– vascular organization was associated with distinct immune cell positioning and activation patterns, and on-chip immune phenotypes correlated with histological joint inflammation scores. We then tested infliximab, tocilizumab, and tofacitinib in parallel and integrated cytokine secretion, extracellular matrix remodeling, immune cell survival and immune cell recruitment into a composite treatment response score. On-chip scores for clinically matched treatments correlated with clinical improvement measured by ΔDAS28–ESR (Pearson r = 0.844). To investigate the cellular basis of heterogeneous drug responses, single-cell RNA sequencing identified SFRP1⁺ remodeling FLS that persisted under treatment and exhibited high inflammatory programs in poorly responding chips. Cell–cell communication analysis linked reduced FN1–CD44 signaling toward SFRP1⁺ remodeling FLS to higher NF-κB activity and drug resistance. We tested the functional relevance of this finding in chips derived from an independent TNFi-nonresponsive patient, where fibronectin supplementation reduced IL-6 secretion and restored the composite treatment response. Together, these findings support patient-derived synovium-on-a-chip as a platform for individualized drug evaluation and identify ECM-mediated niche signaling as a potential therapeutic target for overcoming drug resistance in RA.

**One Sentence Summary:** A patient-derived immunocompetent synovium-on-a-chip recapitulates rheumatoid arthritis joint biology in vitro, accurately tests patient therapeutic response and reveals a SFRP1⁺ FLS-mediated resistance mechanism in rheumatoid arthritis.

## INTRODUCTION

Rheumatoid arthritis (RA) is a systemic inflammatory disease characterized by chronic inflammatory arthritis and progressive joint destruction, affecting approximately 1% of the global population(*1–3*). Delay in effective treatment is associated with increased morbidity and disability. Achieving remission within the first three months remains the strongest predictor of long-term disease control (*4–7*). Conventional synthetic disease-modifying antirheumatic drugs (csDMARDs, e.g., methotrexate) are universally recommended by guidelines as the standard first-line treatment for rheumatoid arthritis, because starting with a biologic DMARD in a treatment-naïve patient increases healthcare costs. The majority (seventy-five to eighty-five percent) do not respond sufficiently to csDMARD monotherapy (*8*). Biologic and targeted synthetic DMARDs (b/tsDMARDs) are approved for treatment of rheumatoid arthritis. Yet, even beyond the csDMARD stage, the treatment strategy is one of trial-and-error, as no reliable method exists to predict which agent will benefit a given patient (*9, 10*). During the typical 3 to 6 months treatment trial, responses are variable, with approximately 30% not responding and 30% responding only partially to b/tsDMARDs (*8, 11, 12*). Thus, active disease persists in a substantial percentage of patients even after 6 to 12 months of treatment, during which time damage accrues.

Recent bulk and single-cell transcriptomic studies of RA synovial tissue and peripheral blood have identified candidate pathogenic populations, including CD90⁺HLA-DRA^hi^ and FAP⁺ fibroblast-like synoviocytes (FLS) (*13, 14*) and IL1B⁺/SPP1^+^ macrophages (*15, 16*). These studies have also proposed stratification strategies based on synovial cellular composition (*17–19*), endotype-specific biomarkers (*20, 21*), or circulating cytokine profiles (*22, 23*) and suggest that these patterns could be helpful in predicting therapeutic response. However, none of these have been translated into clinically actionable assays capable of guiding treatment selection for individual patients. An experimental platform that enables systematic evaluation of drug-specific effects within a patient-specific biological context remains a critical unmet need.

In April of 2025, The U.S. Food and Drug Administration announced plans to replace animal testing requirements for new drug approvals, prioritizing New Approach Methodologies including organ-on-chip technologies (*24*). Existing *in vitro* models of the RA synovium face several key limitations. Simple two-dimensional monolayer cultures of FLS, while widely used, lack the cellular complexity and spatial organization of the native synovial microenvironment. Three-dimensional spheroid and organoid systems incorporating FLS with macrophages or endothelial cells have improved multicellular complexity (*25, 26*), but these static models cannot recapitulate perfusable vasculature or immune cell trafficking. Recent synovium-on-a-chip platforms have begun to address this gap by integrating endothelial channels with FLS under flow conditions, and emerging designs have achieved preliminary lining-sublining compartmentalization (*27, 28*). However, these platforms have not used patient material and instead rely on immortalized cell lines or non-autologous immune sources such as THP-1-derived macrophages, precluding patient-specific findings that would address personalized medicine. This limitation is particularly relevant to autoimmune diseases, in which patient-intrinsic immune dysregulation is central to disease pathogenesis. The absence of autologous, patient-matched immune cells further limits the capacity to capture clinically relevant patient heterogeneity, endotype-specific stromal–immune organization, and individualized drug responses.

In this study, we developed a bioengineered, vascularized, immunocompetent Synovium-on-a-Chip platform that reconstructs the microanatomic organization of human synovium *in vitro*. We enrolled 10 RA patients spanning three synovial tissue endotypes (*29*) and integrated patient-derived FLS from synovial tissue with matched peripheral blood mononuclear cells within a compartmentalized and vascularized microfluidic device. The resulting on-chip synovial niche recapitulated patient-specific endotype identity and revealed that endotype-specific stromal-vascular architecture coordinates immune cell positioning, activation, and survival. Multimodal drug profiling using the Synovium-on-a-Chip platform mirrored clinical treatment outcomes and uncovered a fibronectin 1 (FN1) - CD44-mediated drug resistance mechanism sustained by SFRP1⁺ FLS, enabling systematic interrogation of individualized therapeutic responses and providing a functional platform for personalized treatment selection in RA.

## RESULTS

### Bioengineering a patient-derived, immunocompetent RA synovium-on-a-chip

Anatomically, RA synovium comprises layers, including a lining layer (LL) populated by macrophage-like synoviocytes and FLS, and a deeper sublining layer (SL) containing immune infiltrates and vasculature. An activated RA niche is characterized by FLS hyperproliferation (*30*), pathogenic fibroblast subsets (*17, 31, 32*), leukocyte infiltration (*33*), pro-inflammatory myeloid polarization (*34*), and robust local cytokine production (*35*). To establish a patient-relevant platform, we established an *ex vivo* vascularized, immunocompetent RA synovium-on-a-chip using FLS isolated from patient synovial tissue and matched peripheral blood T and B lymphocytes and monocytes. (**Fig. 1A**).

**Fig. 1.**
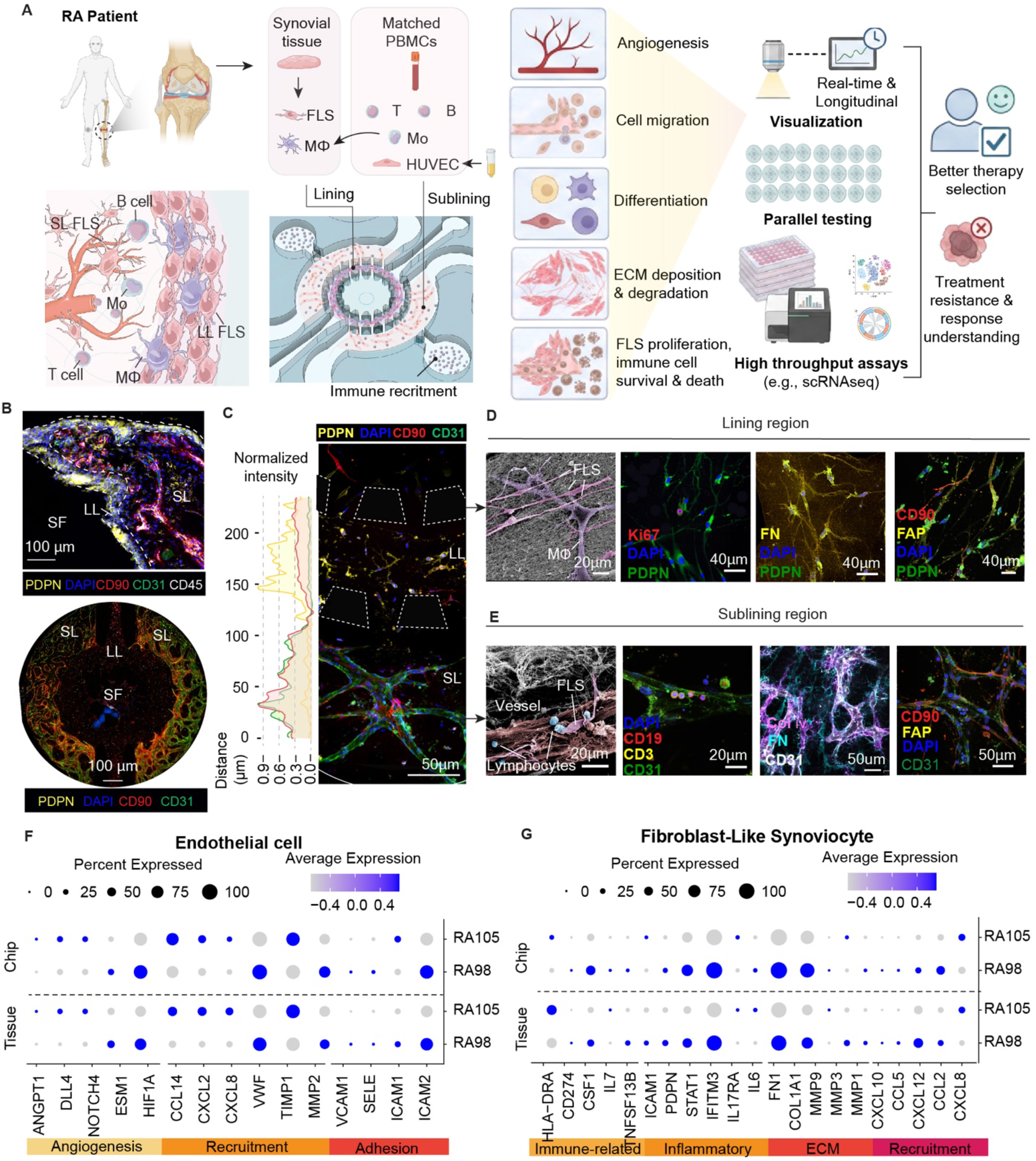
Engineering a patient-derived, immunocompetent rheumatoid arthritis synovium-on-a-chip platform. **(A)** Schematic illustration of the patient-derived RA synovium-on-a-chip platform. Synovial FLS isolated from RA synovial tissue and matched PBMCs are integrated into a compartmentalized microfluidic device that recapitulates the lining and sublining regions of synovium, with side immune cell recruitment channels. The platform enables real-time visualization of angiogenesis and cell migration, parallel testing of serial chips derived from the same patient, and analyses using high throughput assays. **(B)** Representative immunofluorescence images of RA synovial tissue (top) and RA synovium-on-a-chip (bottom) highlighting the spatial organization of the lining layer (LL), sublining layer (SL) and synovial fluid (SF) region. Representative tissue immunofluorescence images are shown from analyses of three tissue sections. **(C)** Spatial distribution of immune and stromal markers across the lining and sublining regions on chip at day 8 (mature niche), showing compartment-specific localization of CD31⁺ endothelial cells, CD90^+^ sublining FLS predominantly in the sublining region, and PDPN⁺ lining FLS enriched at the lining layer. Distance is measured from the outer surface of the sublining region to the inner surface of the lining micropillar array. Fluorescence intensities were independently min–max normalized to 0–1 for each marker. **(D)** Electron microscopic (EM) and immunofluorescence characterization of the lining region in RA synovium-on-a-chip, showing Ki67⁺ proliferating lining FLS, the presence of macrophages within the lining region, extracellular matrix deposition, and activated fibroblast subsets characterized by FAP⁺CD90⁻ expression. **(E)** EM and immunofluorescence characterization of the sublining region, showing perivascular growth of CD90⁺ FLS, infiltration of immune cells, extracellular matrix (ECM) deposition, and activated fibroblast subsets expressing FAP⁺CD90⁺. **(F-G)** Single-cell RNA sequencing dot plots comparing selected gene expression profiles in endothelial cells **(F)** and FLS **(G)** from chips and matched synovial tissue from two patients representing high-and low-inflammatory synovial phenotypes (RA98 and RA105, respectively). Genes are organized into the indicated functional categories. Dot size represents the percentage of cells expressing each gene. Chip samples were collected on day 8. Images in B–E are representative of 3 independent chips. Scale bars are indicated in the images.

The function of pathogenic FLS and immune cells is closely tied to their spatial localization within the synovium (*2, 36*). Recapitulating the spatial segregation of lining and sublining FLS populations is therefore critical for faithfully reproducing RA synovial biology. To replicate this tissue architecture *in vitro*, the microfluidic device was partitioned by arrays of micropillars into three concentric regions corresponding to synovial fluid, lining, and sublining compartments, along with lateral channels for quantifying immune recruitment capacity (**Fig. 1A, Supplemental Fig. 1A, B**). Multiple chips derived from the same patient were used in parallel to allow different assays, replicates, and sample collection at different time points. During chip loading, FLS isolated from RA patient synovial tissue and peripheral blood monocyte-derived macrophages were embedded in the lining region, while CD3⁺ T cells, CD19⁺ B cells, monocytes from peripheral blood, and vascular cells (HUVEC, C2519A) were incorporated into the sublining region within a 3D biomimetic fibrin hydrogel. Over 8 days of culture, FLS proliferated, migrated, and spontaneously adopted spatially distinct phenotypes. FLS localizing proximal to the vasculature progressively upregulated CD90 (sublining FLS marker, **Supplemental Fig. 1C**), whereas FLS more distant from vessels retained high expression of the lining marker PDPN, recapitulating the lining–sublining segregation observed in patient synovial tissue (**Fig. 1B)**. Quantitative spatial analysis further showed that the CD90:PDPN expression ratio decreased with increasing distance from the nearest vascular cell, consistent with in vivo observations (*37*) (**Fig. 1C, Supplemental Fig. 1C, 1E**). This spatially organized differentiation was absent in FLS monoculture, where cells expressed equivalent levels of CD90 and PRG4 (lining FLS marker) without directional phenotypic bias (**Supplemental Fig. 1D, 1E**), highlighting the advantage of the vascularized synovial chip in reconstituting synovial spatial organization. Further characterization of the chip compartments confirmed layer-specific pathological features. In the lining region, scanning electron microscopy (SEM) revealed direct FLS–macrophage contacts, and immunofluorescence demonstrated Ki67⁺ FLS proliferation, ECM deposition, and a pathogenic FAP⁺CD90⁻ fibroblast subset (**Fig. 1D**). In the sublining layer, perivascular CD90⁺ FLS accumulation, immune cell infiltration, ECM remodeling, and FAP⁺CD90⁺ activated fibroblasts were observed (**Fig. 1E**), recapitulating the invasive FAP⁺ fibroblast phenotypes associated with inflammation and joint destruction in RA (*14*).

At the transcriptional level, scRNA-seq of cells dissociated from synovial tissue and cells extracted from synovium-on-a-chip derived from the same synovial tissue demonstrated comparable FLS gene expression profiles across immune-related, inflammatory, ECM, and recruitment-associated programs in both high (RA98) and low (RA105) inflammatory endotypes, with patient-specific inter-individual differences preserved on chip (**Fig. 1G).** Transcriptional correspondence extended to the endothelial and immune compartments. Notably, although all chips were vascularized using HUVECs from the same healthy donor, chip-derived endothelial cells acquired patient-specific transcriptional programs that mirrored synovial endothelial cells (**Fig. 1F**), which suggests that the patient-derived synovial microenvironment actively shapes endothelial identity to adopt disease-relevant transcriptional states despite their distinct donor origins. Endotype-associated immune cell states were also maintained between chip and tissue (**Supplemental Fig. 2C**). Collectively, these findings confirm that the platform recapitulates not only the structural, cellular and transcriptional features of primary RA synovium, but also has the capacity to interrogate person-specific cellular states enabling personalized medicine.

To assess the functional fidelity of the platform, we compared synovium-on-a-chips derived from RA patients with synovium-on-a-chips derived from osteoarthritis (OA) patients. RA chips exhibited a higher proportion of Ki67⁺ FLS than OA chips, recapitulating the hyperproliferative stromal phenotype of RA (**Fig. 2D**). Because FLS actively participate in both ECM synthesis and degradation including fibronectin and collagen I/VI, changes in these matrices may reflect both altered deposition and degradation. To minimize this confounding effect and more specifically assess ECM degradation, we quantified vascular basement membrane–associated collagen IV surrounding CD31⁺ vascular that deposited by a common endothelial cell source across conditions, thereby providing a standardized ECM reference. Collagen IV intensity progressively decreased between Days 3 and 9 in RA chips but increased in OA controls, consistent with enhanced matrix loss and impaired basement-membrane maintenance in the RA microenvironment (**Fig. 2B**). MMP and TIMP profiling also revealed higher secretion of MMP-1 and MMP-3 in RA chips, whereas OA chips secreted more homeostasis-associated MMP-2. In OA chips, relatively low MMP-1 and MMP-3 secretion alongside abundant TIMP-1 and TIMP-2 was consistent with matrix preservation and progressive collagen IV accumulation. In contrast, RA chips exhibited increased MMP-1 and MMP-3 secretion and progressive collagen IV loss despite abundant TIMP secretion, suggesting active stromal remodeling accompanied by net matrix loss **(Fig. 2C)**.

**Fig. 2.**
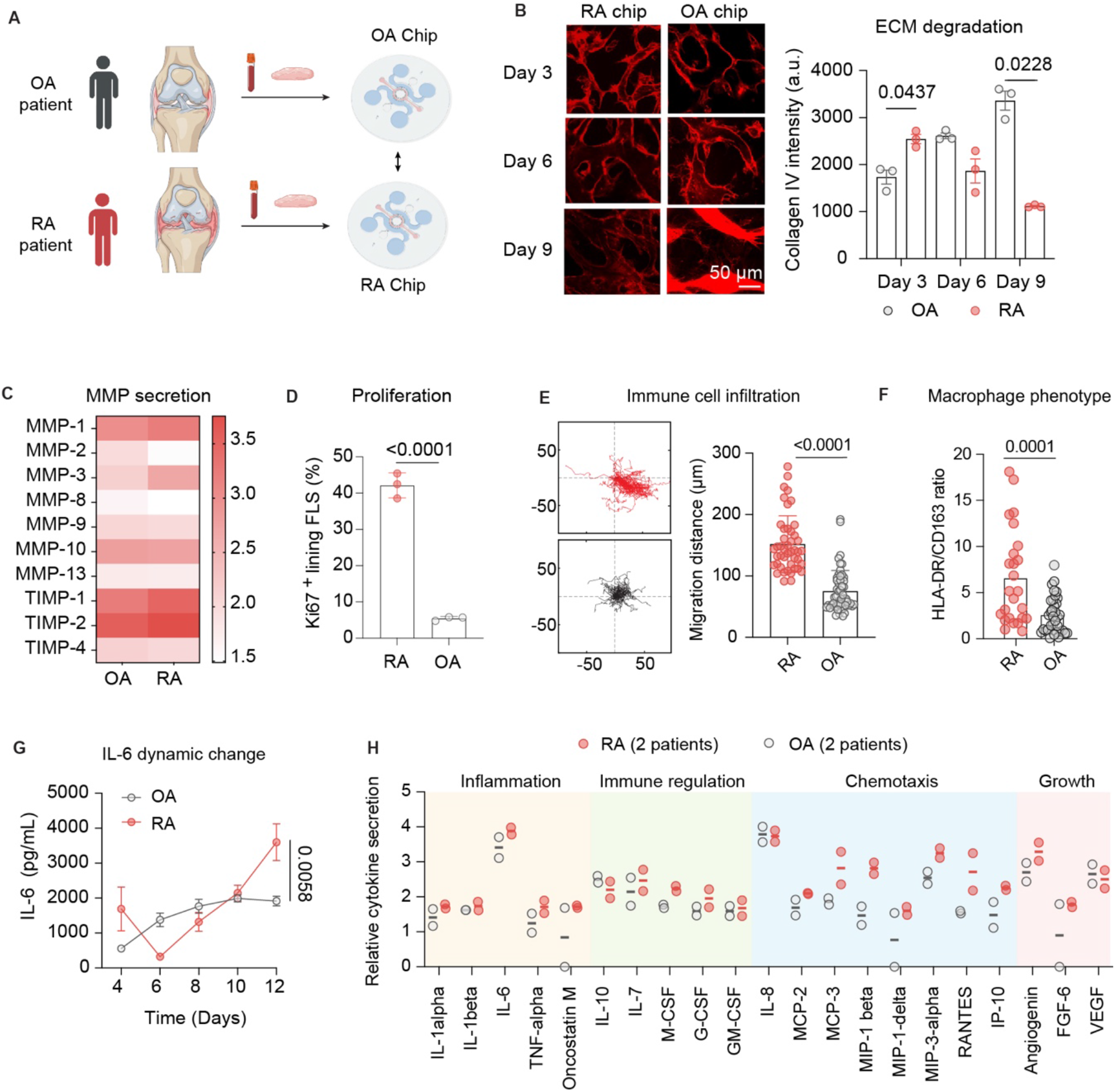
Patient-derived RA synovium-on-a-chip recapitulates key pathological features and the inflammatory microenvironment of rheumatoid arthritis. **(A)** Schematic of the generation of synovium-on-a-chip models using synovial tissue and blood samples from patients with RA or osteoarthritis (OA) for comparative assessment of disease-associated phenotypes. (B) Representative immunofluorescence images and quantification of collagen IV deposition in RA and OA chips on days 3, 6, and 9 (n = 3 chips per condition per time point; Comparisons between RA and OA chips at each time point were performed using two-tailed unpaired Welch’s t-tests with Šídák correction for three comparisons; P values indicated). Scale bar, 50 μm. (C) Heatmap showing secretion profiles of matrix metalloproteinases (MMPs) and tissue inhibitors of metalloproteinases (TIMPs) in OA and RA chip culture medium pooled from 3 independently cultured chips on day 12. Color scale indicates log₁₀-transformed relative secretion values. **(D)** Quantification of Ki67⁺ lining FLS. Each dot represents the mean value for one chip (n = 3 chips; two-tailed unpaired t-test; P < 0.0001). **(E)** Representative immune cell migration trajectories and quantification of migration distance in RA and OA chips (three technical replicates with >45 immune cells per condition, two-tailed unpaired t-test, P <0.0001) **(F)** Quantification of the HLA-DR/CD163 ratio as an indicator of macrophage polarization (n = 3 chips per condition). **(G)** Temporal profiles of IL-6 concentrations in culture medium collected from RA and OA chips on days 4, 6, 8, 10, and 12 (n = 3 chips per condition per time point). IL-6 concentrations on day 12 were compared between RA and OA chips using a two-tailed unpaired t-test (P = 0.0058). **(H)** Multiplex profiling of cytokines associated with inflammation, immune regulation, chemotaxis, and growth in RA and OA chips on day 12. Each dot represents one patient, measured using culture medium pooled from 3 independently cultured chips; 2 patients were analyzed per group. Horizontal lines indicate group means. Values represent log₁₀-transformed relative cytokine secretion. Bars and error bars in (B), (D–G) indicate mean ± SD.

To compare immune cell recruitment and migration dynamics across patient-derived chips, T cells, B cells, and monocytes from the same healthy donor were introduced into the side channels on Day 12, the final day of chip culture. These cells served as a standardized source of recruited immune cells for the endpoint migration assay, while the resident immune compartment was established using patient-matched cells throughout the preceding culture period. RA chips supported substantially greater immune cell migration, consistent with enhanced immune cell recruitment capacity (**Fig. 2E**). Following niche maturation, RA chips also exhibited a higher M1/M2 macrophage ratio, consistent with a shift toward proinflammatory macrophage polarization (**Fig. 2F**). Soluble mediator profiling further revealed distinct inflammatory dynamics. IL-6 secretion rose sharply in RA chips by Day 12, whereas OA chips maintained a comparatively lower and more stable secretion (**Fig. 2G**). Multiplex profiling at Day 12 demonstrated relatively higher secretion of several inflammatory and immune-associated mediators in RA chips, including IL-6, MCP-3, MIP-1β, MIP-3alpha, RANTES, as well as angiogenin (**Fig. 2H**). Collectively, these orthogonal readouts demonstrate that the platform distinguishes RA from OA across stromal proliferation, matrix homeostasis, immune recruitment and polarization, and cytokine output, supporting its disease-specific functional fidelity.

### Endotype-specific stromal–vascular architecture shapes immune cell positioning in the RA synovial niche

We stratified patients into low, mixed, and high inflammatory synovial endotypes using a machine-learning algorithm that incorporates the Hospital for Special Surgery synovitis score and gene expression data as previously published by Orange et al. (*29*). An expert musculoskeletal pathologist semiquantitatively scored 20 histological features from an H&E-stained section of the synovial tissue, integrating the presence or absence of neutrophils, detritus, fibrin, binucleate plasma cells, Russel bodies, and synovial giant cells and the amount of mucoid change, lymphocytic inflammation, lining hyperplasia and plasma cell infiltrate. Patients with a joint inflammation score (JIS) of <3, 3–7, or >7 were classified as low, mixed, or high inflammatory endotypes, respectively (**Fig. 3A**). When examining the engineered synovial niche across synovial endotypes, immunofluorescence imaging revealed distinct immune cell localization patterns. In low inflammatory (low-) endotype chips, CD45⁺ immune cells were predominantly retained within CD31⁺ vascular structures, whereas in high inflammatory (high-) endotype chips, immune cells extravasated from vessels and accumulated in the surrounding tissue. Quantitative analysis showed that low-endotype chips contained a higher overall number of CD45⁺ cells but a smaller fraction outside the vasculature, whereas high-endotype chips contained fewer CD45⁺ cells overall but a significantly greater tissue-localized proportion (P = 0.0353 and P = 0.0363, respectively; **Fig. 3B**). This divergence suggested that inflammatory endotypes differ not only in immune cell abundance but also in the spatial positioning of immune cells relative to the vascular and stromal compartments.

**Fig. 3.**
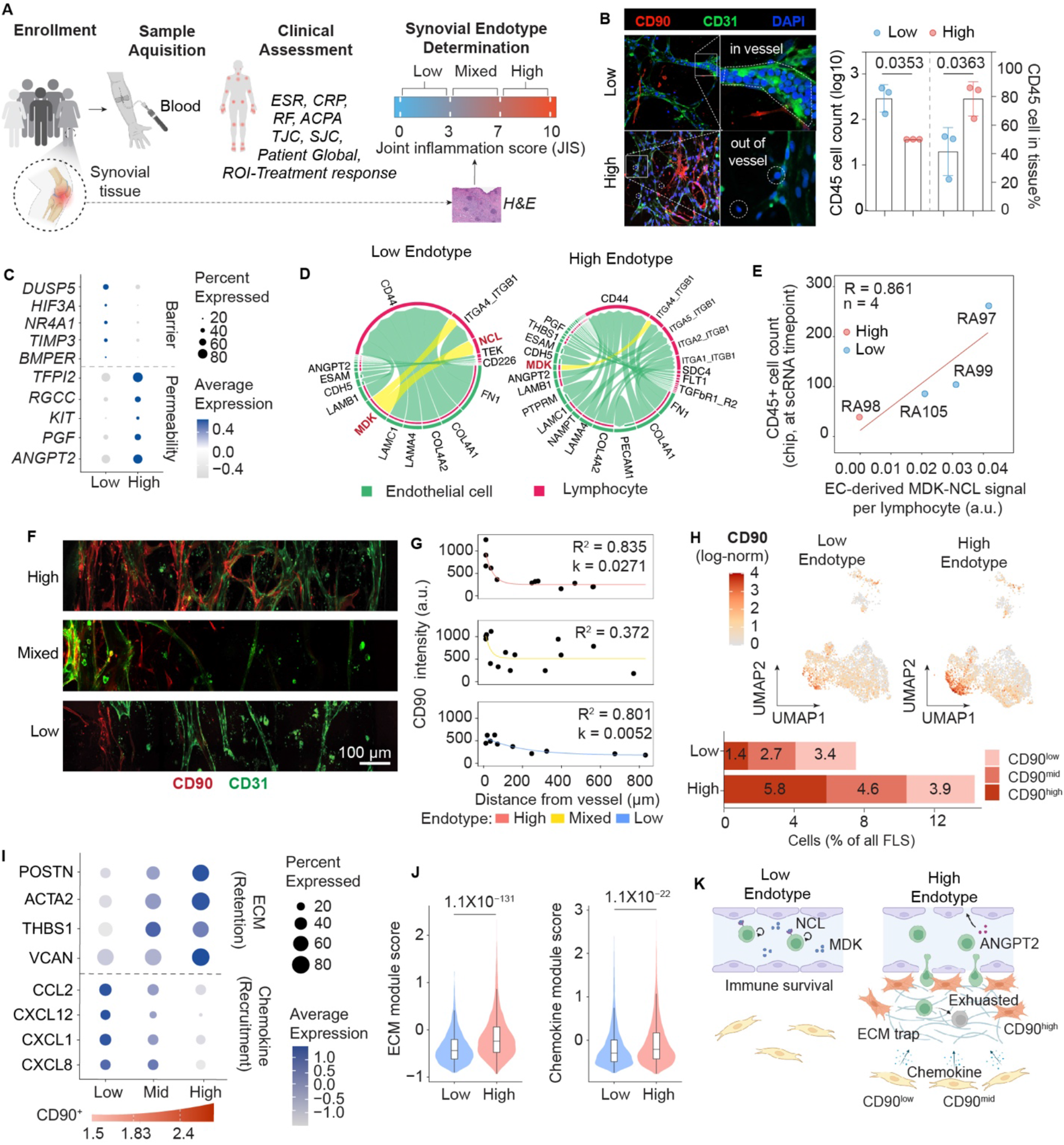
Endotype-specific stromal–vascular architecture shapes immune cell positioning in the RA synovial niche. **(A)** Workflow including patient enrollment, biosample collection, clinical assessment and histology-based synovial inflammatory endotype classification. Clinical assessments included measurements of erythrocyte sedimentation rate (ESR), C-reactive protein (CRP), rheumatoid factor (RF), anti-citrullinated protein antibody (ACPA), tender joint count (TJC), swollen joint count (SJC), and Patient Global Health. Participants permitted record retrieval for collection of treatment response data (ROI, Release of Information). Synovial tissue was evaluated by H&E staining and then stratified into low, mixed, or high inflammatory endotypes according to the joint inflammation score (JIS), as described in the text. **(B)** Representative immunofluorescence images and quantification of endotype-specific immune cell localization relative to CD31⁺ vascular structures. Low-endotype chips contained more CD45⁺ cells but a lower proportion of tissue-localized cells. High-endotype chips showed a higher proportion of CD45⁺ cells outside the vascular compartment and lower number of CD45⁺ cells overall (n = 3 independent patients per endotype; each dot represents the mean of three chips generated from one patient. Bars show mean ± SD across patients. CD45⁺ cell counts were compared using a two-tailed unpaired Welch’s t-test on log10-transformed counts, and the percentages of CD45⁺ cells outside the vascular compartment were compared using a two-tailed unpaired Student’s t-test. P values are indicated.) **(C)** Dot plot showing endothelial expression of barrier- and permeability-associated genes in low- and high-endotype chips. Dot size represents the percentage of cells expressing each gene, and color indicates scaled average expression. **(D)** CellChat analysis of endothelial cell–lymphocyte communication in low- and high-endotype chips. Chord width represents the inferred interaction strength. **(E)** Correlation between endothelial cell-derived MDK–NCL signaling strength per lymphocyte and on-chip CD45⁺ cell count at the scRNA-seq collection time point. (n = 4 patients; each dot represents one patient; Pearson R = 0.861). **(F)** Representative immunofluorescence images showing the spatial organization of CD90⁺ FLS relative to CD31⁺ vessels in high, mixed, and low inflammatory endotypes (representative of n = 3 patients per endotype; images shown from one representative patient per endotype). Scale bar, 100 μm. **(G)** Quantification of CD90 intensity as a function of distance from the nearest CD31⁺ vessel. High-endotype chips exhibited a strong perivascular CD90 gradient, whereas mixed- and low-endotype chips showed weaker spatial gradients. Lines indicate nonlinear decay fits; corresponding R² values and decay constants (k) are shown. **(H)** UMAP feature plots showing CD90 expression in FLS from low- and high-endotype chips, with the distribution of CD90^low^, CD90^mid^, and CD90^high^ states expressed as percentages of all FLS. **(I)** Dot plot showing functional gene programs across CD90^low^, CD90^mid^, and CD90^high^ FLS populations. CD90^high^ FLS were enriched for ECM-associated genes, including *POSTN*, *ACTA2*, and *VCAN*, whereas CD90^low^ and CD90^mid^ FLS expressed chemokine and immune recruitment-associated genes, including *CCL2*, *CXCL12*, *CXCL1*, and *CXCL8*. Dot size represents the percentage of cells expressing each gene, and color indicates scaled average expression. **(J)** Comparison of ECM-retention and chemokine-recruitment module scores between low- and high-endotype FLS, showing significantly increased activity of both programs in the high endotype (P values indicated). For **(H–J)**, scRNA-seq data were obtained from n = 1 patients per endotype. **(K)** Working model of endotype-specific stromal– vascular regulation of immune cell positioning. Low-endotype chips maintain immune cells within an intact vascular niche supported by endothelial MDK–NCL survival signaling. High-endotype chips exhibit ANGPT2-associated vascular leakage and immune cell extravasation, accompanied by chemokine-mediated recruitment by CD90^low/mid^ FLS, an ECM-rich perivascular retention niche formed by CD90^high^ FLS, and exhausted immune cell states.

As the distinct immune cell localization patterns were closely associated with vascular structures, we next examined whether endotype-specific endothelial programs could explain differences in immune retention and extravasation. We retrieved endothelial cells from the chips and analyzed by scRNA-seq. ECs from low-endotype chips expressed genes associated with vascular quiescence and barrier maintenance, including *DUSP5*, *HIF3A*, *NR4A1*, *TIMP3*, and *BMPER*, whereas ECs from high-endotype chips upregulated permeability-and activation-associated genes, including *TFPI2*, *RGCC*, *KIT*, *PGF*, and *ANGPT2* (**Fig. 3C**). CellChat analysis further revealed endotype-specific endothelial-lymphocyte interactions: low-endotype chips showed enriched MDK-NCL signaling, a pathway associated with lymphocyte survival, whereas high-endotype chips exhibited increased permeability-and adhesion-associated communication (**Fig. 3D**). Cross patient analysis also showed that endothelial-derived MDK–NCL interaction strength per lymphocyte was positively associated with on-chip CD45⁺ cell abundance (Pearson R = 0.861, n = 4; **Fig. 3E**), supporting a relationship between endothelial survival signaling and the intravascular immune retention observed in low-endotype chips.

Although increased vascular permeability may facilitate immune cell extravasation, it does not fully explain why immune cells accumulate within the synovial niche after leaving the vessel. We therefore asked whether stromal spatial organization actively contributes to immune recruitment and retention. Immunofluorescence analysis showed that CD90 expression was strongly associated with vessel proximity, with pronounced perivascular enrichment of CD90⁺ FLS in high-endotype chips and an exponential decrease in CD90 intensity with increasing distance from the vessel (k = 0.0271, R² = 0.835). This spatial gradient was weaker in mixed-endotype chips and largely absent in low-endotype chips (k = 0.0052, R² = 0.801; **Fig. 3F**). To define the functional significance of this spatial organization, we used scRNA-seq to identify CD90⁺ FLS based on log-normalized CD90 expression (>1.5) and further stratified these cells into CD90^low^ (1.5-1.83), CD90^mid^ (1.83-2.4), and CD90^high^ (>2.4) populations according to their relative CD90 expression levels. High-endotype chips contained a greater proportion of CD90⁺ FLS, and a higher proportion of CD90^high^ population CD90^+^ FLS than low-endotype chips (**Fig. 3H**). Further analysis showed that CD90^high^ FLS enriched in the perivascular niche expressed ECM retention genes including *POSTN*, *ACTA2*, *THBS1*, and *VCAN*, whereas CD90^mid^ and CD90^low^ FLS expressed chemokines including *CCL2*, *CXCL12*, *CXCL1*, and *CXCL8*, capable of recruiting immune cells away from the vasculature and into the tissue (**Fig. 3I**). Correlation analysis further confirmed that ECM module scores increased progressively with CD90 expression level among CD90^+^ FLS, whereas chemokine scores showed the inverse pattern (**Supplemental Fig. 3C-D**). GO biological process analysis and module scoring consistently confirmed upregulation of ECM organization and chemokine-mediated signaling pathways in high endotype FLS (**Supplemental Fig. 3E)**. Consistent with the increased proportion of the CD90⁺ FLS in the high endotype, both ECM and chemokine module scores were significantly higher in high-endotype than in low-endotype FLS (**Fig. 3J**), indicating that immune recruitment and stromal retention are coordinated features of the high-endotype niche.

Together, these data support a coordinated stromal-vascular model of immune cell positioning in RA (**Fig. 3K**), in which intact vascular barriers and endothelial MDK-NCL signaling retain immune cells within a protected intravascular niche in low inflammatory synovial endotype RA, activated and permeable vessels permit immune extravasation, while chemokine-producing CD90^low/mid^ FLS recruit immune cells into the tissue and ECM-rich CD90^high^ FLS retain them within the perivascular niche in high inflammatory synovial endotype RA. This spatial organization was consistent with the prominent perivascular accumulation of immune cells observed by hemotoxylin and eosin stain of synovial tissue from the high-inflammatory endotype (**Supplemental Fig. 3F**). Patient synovial endotypes are thus organized through spatial coordination among endothelial programs, FLS subsets, and ECM architecture.

### Stroma-shaped immune behaviors reflect patient-specific synovial inflammation

To comprehensively characterize RA patient-specific states on chip, we profiled three functional compartments: vascular, FLS, and immune. The vascular module included vessel area and collagen IV deposition **(Fig.4A, Supplemental Figure 5)**; the FLS module included immune cell recruitment, IL-6 secretion, and proliferation **(Fig.4A, Supplemental Figure 4)**; and the immune module included CD45⁺ cell count, CD45⁺ proportion within the tissue compartment, and an inflammatory immune phenotype score, defined as the combined proportion of Th17 cells, plasma cells, non-classical monocytes, and M1 macrophages (**Fig. 4A, Supplemental Fig. 4 and 7**). Among the three compartments, the immune module showed the strongest concordance with JIS-defined endotypes (**Fig. 4A**). Because stromal–vascular architecture shaped immune cell survival and residence within the chip, we used immune cell behavior as a functional output of the stromal– vascular-immune programs described above. The immune module captured three distinct dimensions of immune cell behavior: total CD45⁺ cell count reflected immune survival and vascular retention, CD45⁺ proportion within tissue reflected extravasation and tissue infiltration, and the proinflammatory immune phenotype score reflected immune activation within the synovial niche. Quantitative analysis confirmed that these immune features were differentially associated with local tissue inflammation. CD45⁺ cell count negatively correlated with JIS (R = −0.7448, p = 0.0213), whereas CD45⁺ proportion within tissue positively correlated with JIS (R = 0.8333, p = 0.0083). The proinflammatory immune phenotype scores also showed the strong positive association with JIS (R = 0.9167, p = 0.0013; **Fig. 4B**). Thus, high inflammatory endotypes were not characterized by greater total immune abundance, but by a redistribution of immune cells from the vascular compartment into the tissue and by acquisition of proinflammatory phenotypes within the niche microenvironment. Consistent with this interpretation, progressive enrichment of tissue-localized and proinflammatory immune subsets with increasing JIS was accompanied by a corresponding decline in total CD45⁺ cell count and proliferative capacity (**Fig. 4C**). This pattern suggests that the highly inflammatory perivascular niche in high-endotype chips promotes immune extravasation and activation but does not support sustained immune expansion. In contrast, low-endotype chips retain more immune cells within the vascular compartment, consistent with the endothelial survival and barrier programs identified (**Fig. 3**). To determine whether this activation– proliferation trade-off reflects immune dysfunction *in vivo*, we performed scRNA-seq on matched synovial tissue from high and low endotype RA patients. T cells from high-endotype tissue exhibited significantly elevated exhaustion scores alongside higher cytotoxicity (**Fig. 4D**), consistent with prior reports of expanded PD-1⁺ exhausted T cell populations in highly inflamed RA synovium (*2, 33, 38*). These findings suggest that immune cells in high-endotype niches are not simply more abundant, but are spatially recruited, activated, and progressively driven toward functional exhaustion.

**Fig. 4.**
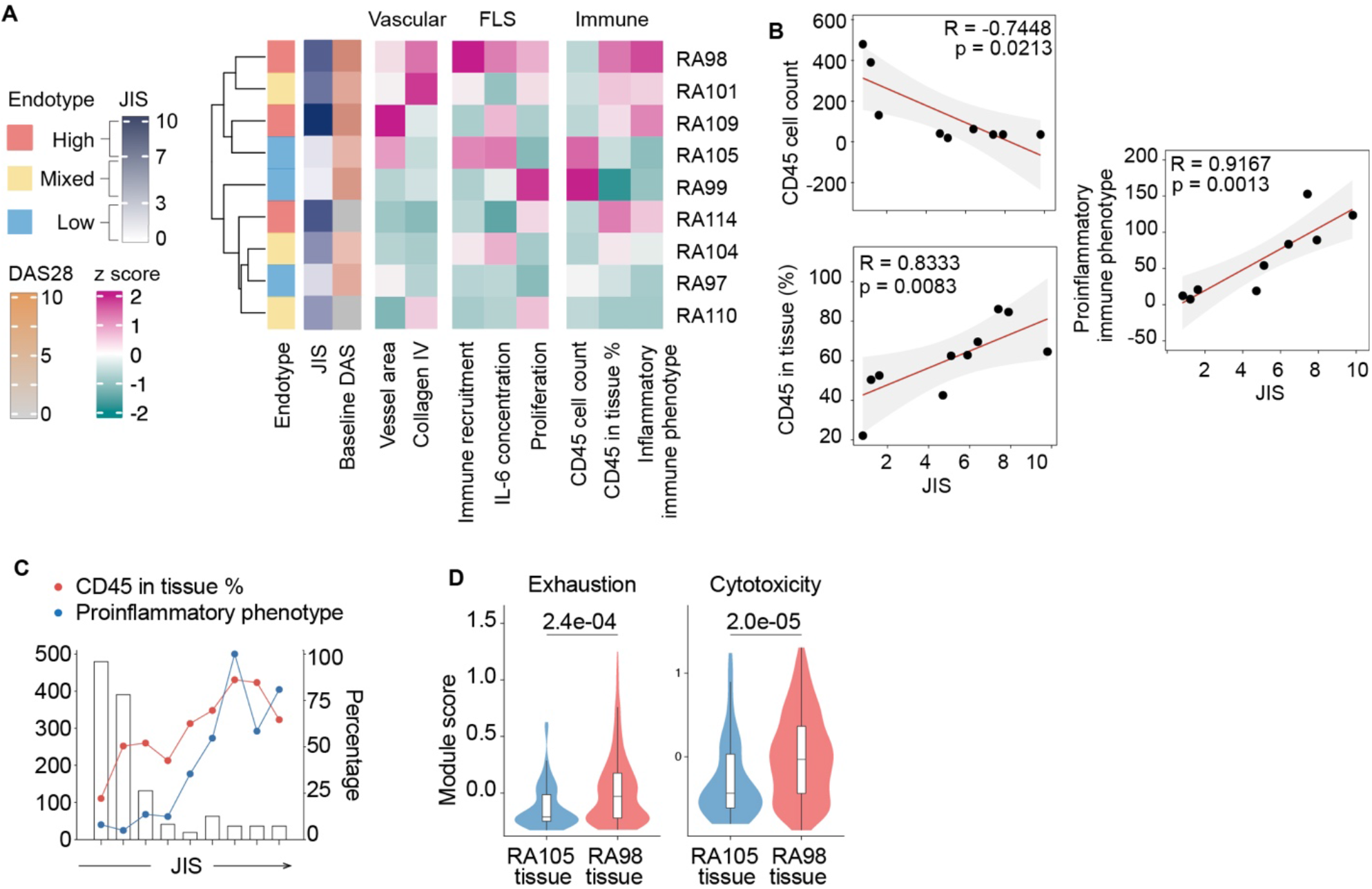
Stromal-shaped immune behaviors reflect patient-specific synovial inflammation. **(A)** Heatmap of multimodal chip readouts across RA patients. Readouts were grouped into vascular, FLS, and immune modules. Each row represents one patient. Values represent the means of 3 chips per patient and were z-score standardized across 9 patients for each readout. Left annotations indicate baseline Disease Activity Score (DAS-28), synovial inflammatory endotype, and JIS. Readouts were collected on day 8, except for immune cell recruitment, which was assessed on day 12. **(B)** Correlation of JIS and on-chip immune readout, CD45⁺ cell count, the percentage of CD45⁺ cells within the tissue compartment, and the proinflammatory immune phenotype score. Each dot represents one patient, using the mean of three chips (n = 9 patients). Spearman correlation coefficients and two-tailed P values are shown. Red lines indicate linear regression fits for visualization; shaded bands indicate 95% confidence interval of the fit. The proinflammatory immune phenotype score was calculated as the sum of four within-lineage percentages: Th17 cells among T cells, plasma cells among B cells, non-classical monocytes among monocytes, and M1 macrophages among macrophages. **(C)** Patient-level CD45⁺ cell counts (bars, left axis), percentages of CD45⁺ cells within the tissue compartment (red points, right axis) and normalized to the maximum patient-level proinflammatory immune phenotype scores (blue points, right axis), ordered by increasing JIS. Each bar or point represents a value derived from the mean of three chips per patient. **(D)** T cell exhaustion and cytotoxicity module scores derived from matched synovial tissue scRNA-seq from RA105 (low endotype; 61 cells) and RA98 (high endotype; 353 cells). Module scores were calculated with Seurat AddModuleScore using exhaustion (*TOX, TIGIT, HAVCR2, LAG3, PDCD1, CTLA4, ENTPD1, IKZF2*) and cytotoxicity (*GZMB, GNLY, NKG7, PRF1, GZMA, GZMH*) gene sets. Violins show the distribution of scores; boxes show the median and interquartile range. P values, two-sided Wilcoxon rank-sum test at the single-cell level; the comparison is between cells from 3-4 tissue sample per patient.

Together, these results demonstrate that stromal–vascular programming of immune cell fate translates into clinically relevant phenotypes. Rather than relying on cross-sectional immune cell abundance alone, the chip captures immune cell localization, survival, and activation as complementary readouts of patient-specific synovial inflammation and systemic disease activity.

### Multimodal on-chip drug profiling reveals drug-specific response signatures and enables personalized therapy selection

Our finding identified immune cell survival and recruitment as functional outputs of stromal– vascular niche organization and linked on-chip immune phenotypes to JIS. Together with the characteristic inflammation and ECM degradation observed in RA, these findings guided the selection of four complementary readouts for on-chip drug profiling: inflammatory cytokine production, ECM remodeling, immune cell survival, and immune cell recruitment **(Fig. 5A)**. RA patients spanning all three inflammatory endotypes were treated on-chip with tocilizumab (IL-6 receptor antagonist, IL6Ra), infliximab (TNF-alpha inhibitor, TNFi), and tofacitinib (Janus kinase inhibitor, JAKi). Patient-specific drug responses and multivariate treatment outcome prediction on the RA synovium-on-a-chip are shown in **Supplemental Figure 6**. Drug-induced changes were expressed as the difference between treated and untreated chips (Δ values). To generate a single quantitative measure of on-chip therapeutic response, the four readouts were each normalized and z-scored across the three treatments and integrated into a composite chip treatment response score, computed as the mean of the four z-scored channels (**Fig. 5C**). The resulting On-chip score captures both the magnitude and breadth of drug-induced changes.

**Fig. 5.**
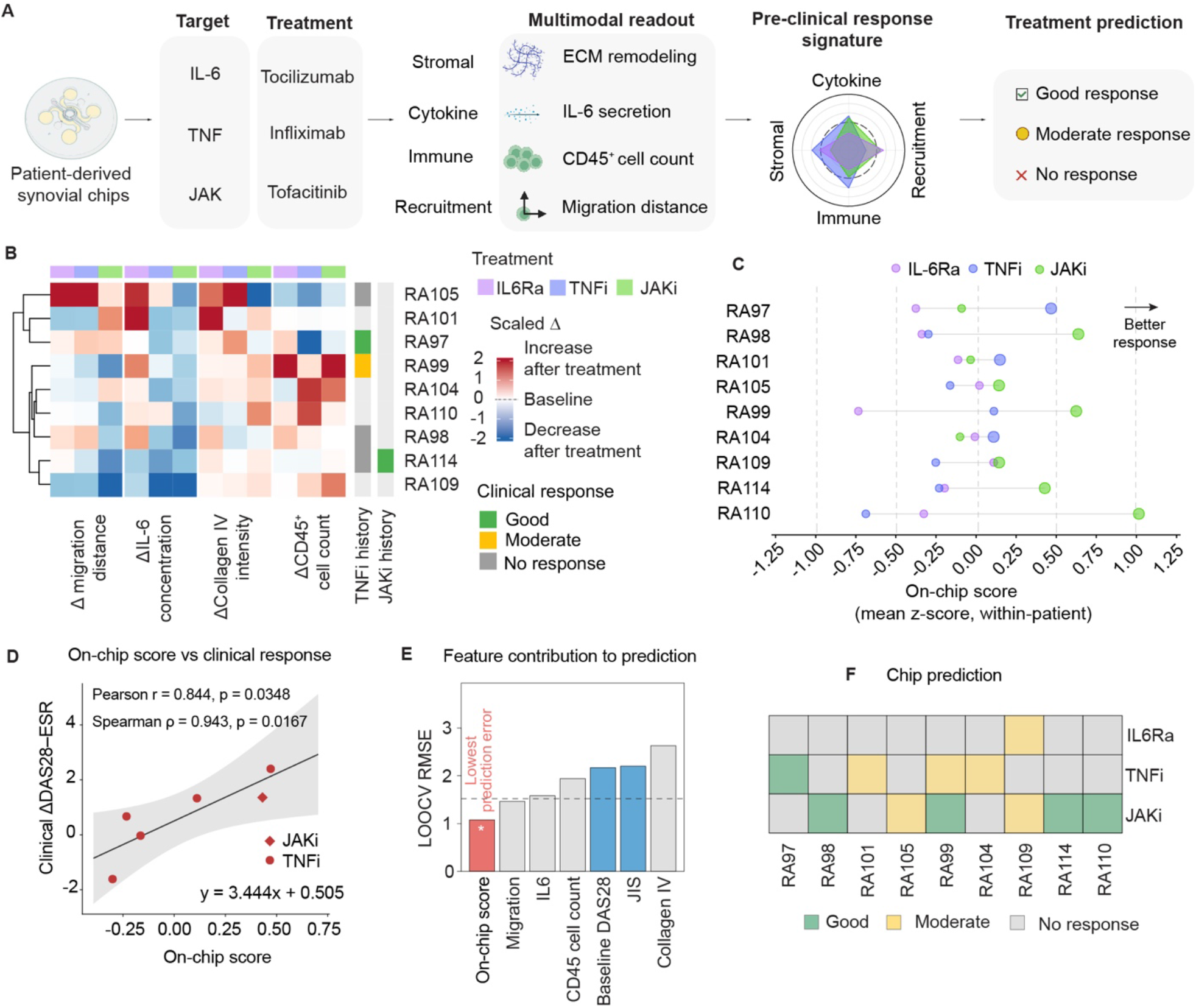
Chip-based multimodal inflammatory signatures predict patient-specific therapeutic response. **(A)** Workflow for treatment prediction using patient-derived RA synovium-on-a-chip. Patient-derived chips were subjected to parallel drug perturbation with tocilizumab (IL-6 receptor antagonist, IL-6Ra), infliximab (TNF inhibitor, TNFi) and tofacitinib (JAK inhibitor, JAKi). Four functional readouts were quantified: stromal (ECM remodeling), cytokine (IL-6 secretion), immune (CD45⁺ cell count) and recruitment (immune cell migration distance). Treatment-induced changes relative to matched untreated controls were calculated as fold changes (treated/untreated), standardized within each patient across the three drug treatments (IL-6Ra, TNFi and JAKi), and combined into a composite on-chip response score to predict therapeutic outcome; the four features are also displayed as radar plots for visualization. **(B)** Heatmap of treatment-induced changes in the four readouts across nine patients and three drug conditions per patient. Rows represent patients, and columns represent readout–treatment combinations. Colors indicate standardized treatment-induced changes, with red indicating increases and blue indicating decreases. Patients were hierarchically clustered using unsupervised Ward’s linkage (ward.D2). Top annotations indicate on-chip treatment; right annotations indicate clinical response history to TNFi and JAKi treatment. Each value represents the mean of 3 chips per condition. **(C)** Composite treatment response score (mean of the four z-score–normalized readouts) quantified for each patient across IL-6Ra, TNFi and JAKi treatments, demonstrating patient-specific and drug-specific variation in composite response magnitude. **(D)** Correlation the composite on-chip response score for the clinically matched treatment and clinical improvement measured as ΔDAS28–ESR. Six matched patient–treatment observations from 5 independent patients are shown. Circles indicate TNFi and diamonds indicate JAKi treatment. Pearson r = 0.844, P = 0.0348; Spearman ρ = 0.943, P = 0.0167; both tests were two-tailed. The line represents the fitted linear regression (y = 3.444x + 0.505), and the shaded band indicates 95% confidence interval of the fit. **(E)** Feature contribution to prediction assessed by LOOCV root mean square error (RMSE) for individual features (migration distance, IL-6, collagen IV and CD45⁺ count, grey), integrated composite score (red) and clinical predictors (baseline DAS28 and JIS-defined inflammatory endotype, blue). The dashed line shows the null model baseline; The integrated composite score achieved the lowest RMSE. The asterisk denotes a prediction error significantly lower than expected by chance for the composite score (permutation test, p = 0.021). **(F)** Chip-based predicted treatment-response categories across nine patients and three drug conditions. Composite on-chip scores were converted into predicted ΔDAS28–ESR values using the regression model in panel D and classified as good, moderate, or no response according to clinical standard (Good: ΔDAS28–ESR>1.2; Moderate: 0.6 <ΔDAS28– ESR ≤ 1.2; No response: ΔDAS28–ESR ≤ 0.6). Green, yellow, and gray indicate the respective predicted categories.

Unsupervised hierarchical clustering of treatment-induced changes (Δ values) revealed heterogeneous response profiles across patients and therapies, shown alongside available clinical treatment-response histories **(Fig. 5B)**. The heatmap also exposed potential mechanisms of drug resistance: in RA98, TNFi treatment reduced IL-6 concentration yet simultaneously enhanced immune cell recruitment, which may contribute to treatment failure. More broadly, IL-6Ra and TNFi increased immune cell recruitment capacity in RA97, RA98, and RA105, while suppressing it in RA101. These divergent patterns indicate that both drug-specific and patient-specific factors govern therapeutic response at a resolution finer than the synovial tissue endotype stratification or single readout can capture alone. This is a unique window into patient-specific differences in biological response to the same treatment. Patients exhibited highly individualized response profiles, with no single therapeutic agent achieving uniformly high scores across all patients.

We next evaluated whether the on-chip score aligned with each patient’s real-world treatment response. In our initial cohort of nine patients tested on chip, one had documented prior clinical treatment with a JAKi and five had documented prior clinical treatment with TNF inhibitors with outcomes ranging from no response to good response (**Fig. 5B, Supplemental Table 1**). We correlated the on-chip treatment response score with their clinically observed ΔDAS28–ESR. The chip treatment response score correlated with clinical disease activity changes with a Pearson R of 0.844 (p = 0.0348; **Fig. 5D**) and a linear relationship of Y = 3.444X + 0.505, linking *ex vivo* multimodal drug effects to *in vivo* therapeutic outcomes. Leave-one-out cross-validation (LOOCV) showed that the integrated composite score achieved the lowest prediction error (LOOCV root mean square error), significantly outperforming each individual chip readout as well as standard clinical predictors (baseline DAS28 and JIS-defined inflammatory endotype; **Fig. 5E**), supporting that multimodal integration predicts therapeutic response more accurately than individual biomarkers or clinical stratification alone.

Applying established EULAR response criteria (Good: ΔDAS28–ESR>1.2; Moderate: 0.6 <ΔDAS28–ESR ≤ 1.2; No response: ΔDAS28–ESR ≤ 0.6) (*39*) to the linear prediction model, we classified each patient’s on-chip composite treatment response to the tested therapeutics as good, moderate, or no response (**Fig. 5F**). The resulting matrix revealed patient-specific therapeutic susceptibility profiles. Although TNFi was the drug class most commonly received clinically in this cohort, chip-based screening predicted that several patients could benefit more from JAKi or IL-6Ra, suggesting that rapid, patient-specific clinical drug screening could enable selection of the optimal therapy within the therapeutic window and potentially achieve disease control before irreversible joint damage occurs. It could also help prioritize the most promising candidate therapy and reduce reliance on sequential empirical treatment selection.

### scRNA-seq of drug-treated synovial chips reveals FLS state-specific mechanisms of therapeutic response and resistance

Even within the low inflammatory endotype, RA97, RA99, and RA105 exhibited markedly different responses to TNFi and JAKi: On-chip, RA97 responded to TNFi but not JAKi, RA99 responded to JAKi and had moderate TNFi sensitivity, and RA105 with no response to TNFi and a moderate response to JAKi (**Fig. 6A**). The patient’s actual real world TNFi treatment response aligned with on-chip treatment TNFi response in all three patients. Because on-chip drug efficacy correlated most strongly with FLS-associated stromal markers, we performed single-cell RNA sequencing (scRNA-seq) on synovium-on-a-chip cultures from these three patients under three conditions (untreated control, TNFi, and JAKi) to investigate the transcriptional basis of drug-specific response heterogeneity within the FLS compartment (**Fig. 6A**). Unsupervised clustering identified eight transcriptionally distinct FLS functional states (**Fig. 6B**). Baseline composition differed markedly across patients: RA105 was dominated by IFN-activated FLS (37%) with a substantial SFRP1⁺ remodeling population (17%), RA97 was enriched in Recruitment lining FLS (27%), and RA99 was dominated by Metabolic FLS (46%) (**Fig. 6C**). These compositional differences, undetectable by conventional histopathological scoring, established distinct stromal baselines from which subsequent drug responses diverged.

**Fig. 6.**
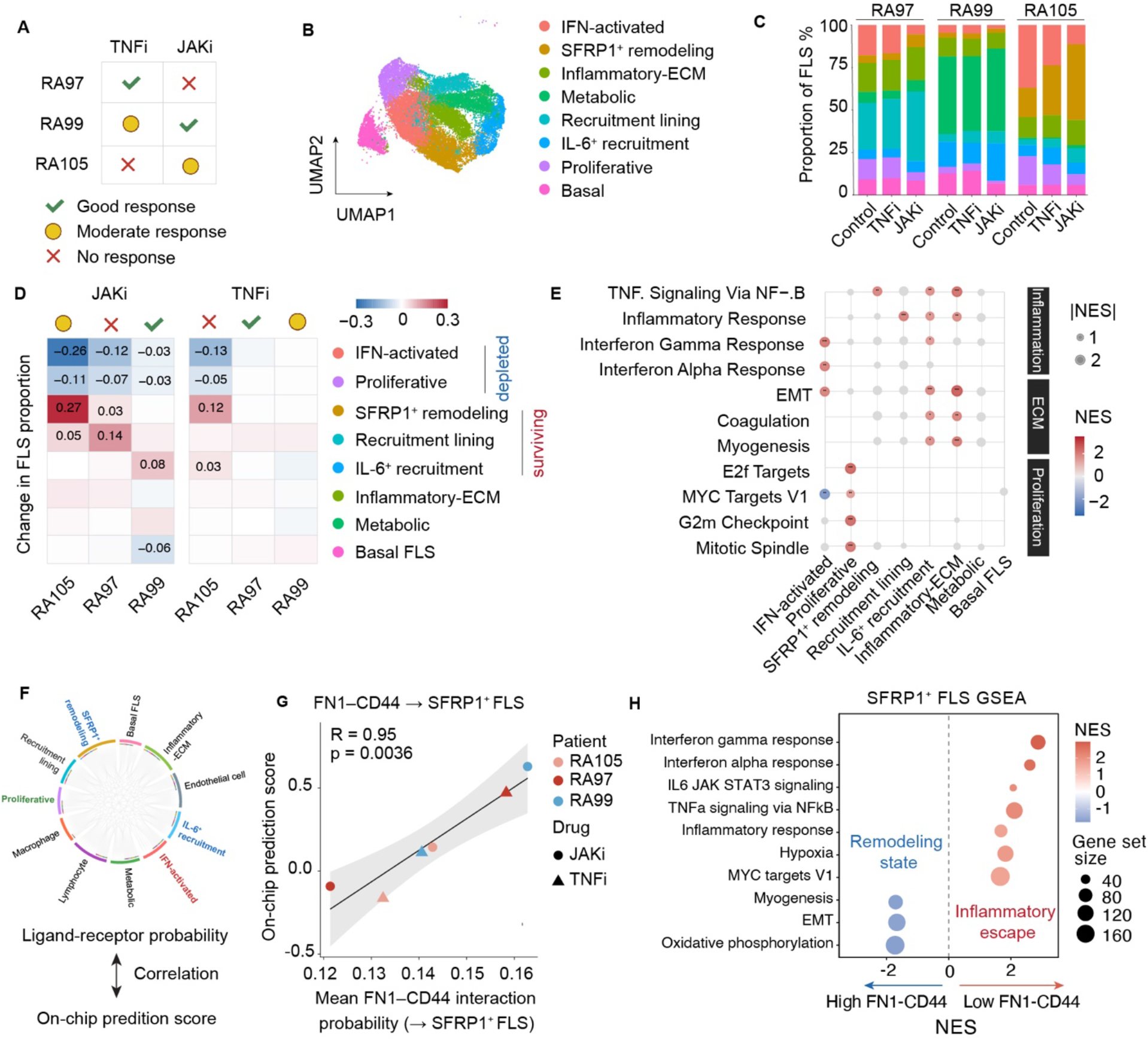
Single-cell transcriptomic analysis reveals FN1-CD44 niche signaling as a determinant of drug response heterogeneity in on-chip RA synovium. **(A)** Summary of chip-predicted drug response profiles for three RA patients form low inflammation endotype (RA97, RA99, RA105) treated with TNFi or JAKi on the synovium-on-a-chip platform. Green check: good response; yellow circle: moderate response; red cross: no response. (B) UMAP projection of scRNA-seq data from FLS isolated across all patients and treatment conditions, colored by eight transcriptionally distinct FLS subpopulations identified by unsupervised clustering. (C) Stacked bar plot showing the proportional composition of eight FLS subpopulations across patients (RA97, RA99, RA105) and treatment conditions (Control-No treatment, TNFi, JAKi). (D) Heatmap of proportion changes for each FLS subpopulation following TNFi or JAKi treatment relative to untreated control, shown for each patient. Subpopulations are grouped into depleted (IFN-activated, Proliferative) and surviving (SFRP1⁺ remodeling, Recruitment lining, IL-6⁺ recruitment) categories. Color scale indicates the magnitude and direction of proportion change. (**E**) Bubble plot of gene set enrichment analysis (GSEA) across eight FLS subpopulations using Hallmark gene sets. Bubble size indicates the absolute normalized enrichment score (|NES|); color indicates NES direction (red, positive enrichment; blue, negative enrichment; grey, nonsignificant enrichment). Pathways are grouped into Inflammation, ECM, and Proliferation categories. (**F**) Schematic of the analytical workflow linking CellChat ligand-receptor interaction analysis to on-chip drug prediction scores. Chord diagram illustrates FN1-CD44 signaling interactions among all cell types on chip, including FLS subpopulations, endothelial cells, lymphocytes, and macrophages. Interaction probabilities for each sender-receiver pair were correlated with on-chip prediction scores across all patient-drug conditions. (**G**) Scatter plot of mean FN1-CD44 interaction probability directed toward SFRP1⁺ FLS versus on-chip prediction score across six patient-drug conditions. Each dot represents one patient-drug condition; dot color indicates patient identity (RA105, RA97, RA99) and dot shape indicates drug (circle, JAKi; triangle, TNFi). Pearson’s correlation coefficient and two-tailed P value are shown (r = 0.9503, P = 0.0036). The solid line indicates the linear regression fit, and gray shading indicates the 95% confidence interval. (**H**) GSEA of SFRP1⁺ FLS transcriptional programs associated with CellChat-inferred FN1–CD44 signaling. Positive NES (red) indicate enrichment associated with low FN1–CD44 signaling, whereas negative NES (blue) indicate enrichment associated with high FN1–CD44 signaling. Bubble size represents gene set size, and color indicates NES. All displayed pathways have adjusted P < 0.05.

Comparison of subpopulation proportions between control and drug-treated conditions revealed a conserved pattern of stromal reorganization in non-responders (**Fig. 6D**). In both JAKi and TNFi non-responder or moderate response conditions, IFN-activated FLS were preferentially depleted (RA105 JAKi: −25%; RA105 TNFi: −13%; RA97 JAKi: −12%), whereas SFRP1⁺ remodeling FLS expanded reciprocally (RA105 JAKi: +27%; RA105 TNFi: +12%; RA97 JAKi: +3%). In contrast, the JAKi responder RA99 exhibited markedly attenuated compositional shifts in both IFN-activated (−2.6%) and Proliferative FLS (−2.58%), suggesting that preservation of these populations reflects effective pathway suppression rather than resistance-associated remodeling.

Gene set enrichment analysis (GSEA) revealed a transcriptional difference between FLS subpopulations that expanded versus those depleted under drug pressure **(Fig. 6E)**. Surviving clusters, SFRP1⁺ remodeling, Recruitment lining, and IL-6⁺ recruitment FLS, were enriched for TNFα/NF-κB signaling, inflammatory response, epithelial–mesenchymal transition (EMT), and coagulation pathways, yet showed minimal enrichment for proliferative programs. In contrast, depleted clusters segregated into two distinct vulnerability profiles: Proliferative FLS were enriched for E2F targets and G2M checkpoint programs, consistent with direct targeting of cycling cells, whereas IFN-activated FLS were enriched for interferon-response signatures, consistent with their selective depletion by JAKi-mediated blockade of JAK-STAT signaling. This transcriptional partitioning indicates that drug-sensitive FLS are eliminated through pathway-specific mechanisms, while resistant subpopulations persist by maintaining inflammatory and tissue-remodeling programs that operate independently of the targeted signaling axis.

To investigate how patient-specific niche contexts regulate the functional behavior of surviving FLS subpopulations, we applied CellChat ligand–receptor interaction analysis across all cell types on chip, including FLS clusters, endothelial cells, lymphocytes, and macrophages and correlated interaction probabilities with on-chip treatment response scores across all six patient– drug conditions. This unbiased screen identified fibronectin 1 (FN1)-CD44 signaling directed from all cell types in the chip toward SFRP1⁺ FLS as the pathway most consistently associated with treatment outcome: FN1-CD44 interaction probabilities were elevated in responder relative to non-responder conditions across all sender cell populations (**Fig. 6F**). The mean FN1-CD44 interaction probability targeting SFRP1⁺ FLS positively correlated with on-chip treatment response scores (Pearson r = 0.9503, P = 0.0036; **Fig. 6G**), establishing the strength of ECM-mediated niche signaling to SFRP1⁺ FLS as a quantitative predictor of treatment outcome.

To determine the transcriptional consequences of FN1-CD44 signaling on SFRP1⁺ FLS identity, we correlated individual gene expression levels in SFRP1⁺ FLS with FN1-CD44 signal strength across conditions (**Fig. 6H, Supplemental Figure 8C**). Under high FN1-CD44 signaling, SFRP1⁺ FLS upregulated quiescence-associated genes including *TIMP3* and *SERPINE2* and were enriched for oxidative phosphorylation and epithelial-mesenchymal transition (EMT) programs. Conversely, under low ECM-CD44 signaling, SFRP1⁺ FLS upregulated interferon-stimulated genes including *IFITM3*, *IFITM2*, and *NNMT*, with enrichment for interferon-gamma response (NES = 2.87), IL-6/JAK-STAT3 signaling (NES = 2.07), and TNFα/NF-κB programs (NES = 2.11). These findings demonstrate that FN1-CD44 niche signaling actively maintains SFRP1⁺ FLS in a quiescent, non-inflammatory state, and that attenuation of this signal licenses their transition toward a pro-inflammatory transcriptional program capable of sustaining synovial inflammation independent of the targeted pathway.

### Reciprocal LXRα and NF-κB activity accompanies FN1-CD44 signaling in SFRP1⁺ FLS

Having established that FN1-CD44 signaling in SFRP1⁺ FLS tracks on-chip treatment response (**Fig. 6**), we asked which transcriptional regulators accompany this signaling. We inferred transcription factor (TF) activity in SFRP1⁺ FLS using DoRothEA and ranked each TF by its correlation with the FN1-CD44 interaction probability across the treated conditions (**Fig. 7A).**

**Fig. 7.**
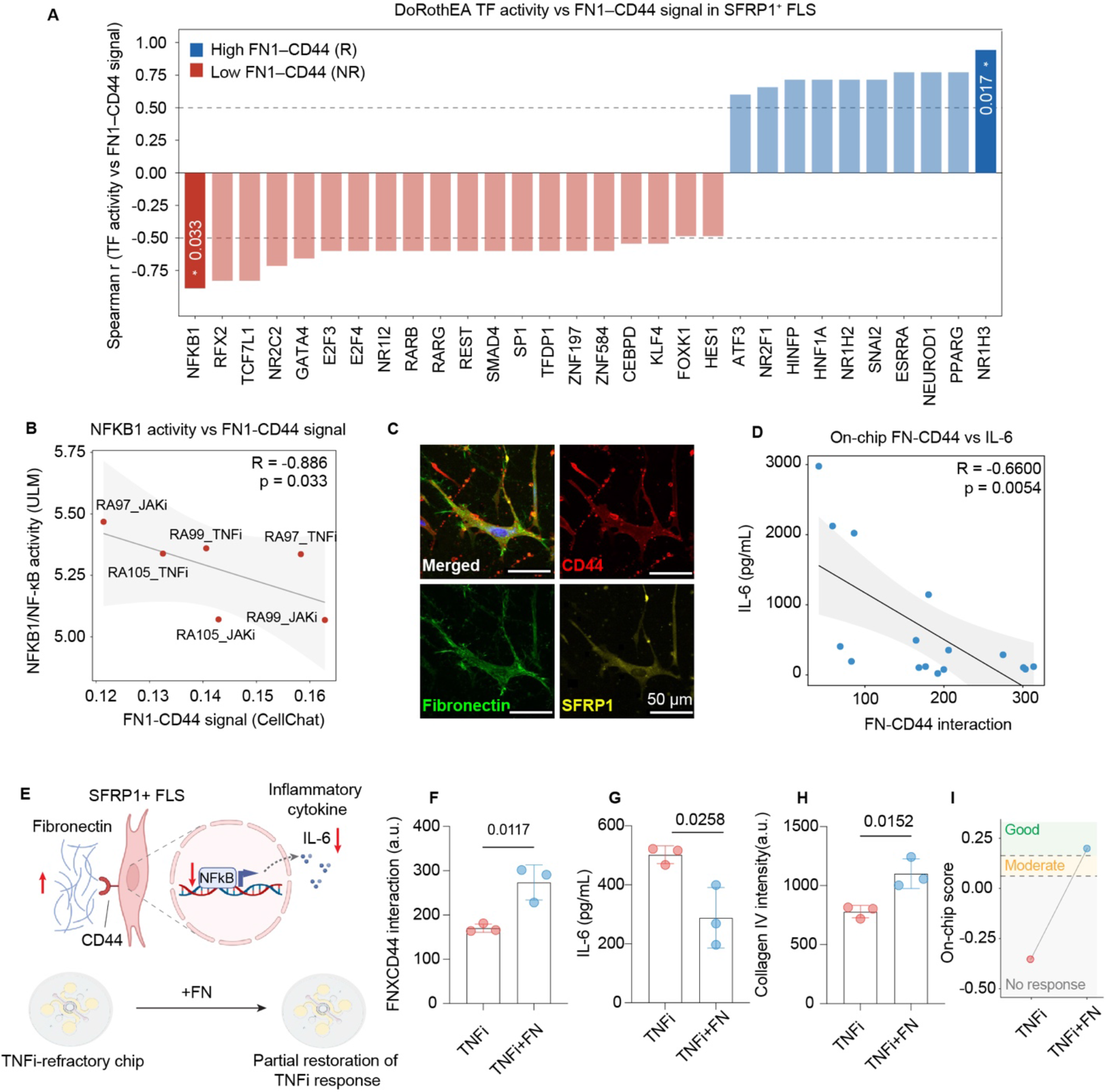
FN1–CD44 niche signaling in SFRP1⁺ FLS is associated with NF-κB activity, and fibronectin supplementation partially restores TNFi responsiveness. **(A)** Spearman correlation analysis between DoRothEA-inferred transcription factor activity and CellChat-derived FN1– CD44 interaction probability in SFRP1⁺ FLS across six treated patient–drug conditions. Transcription factors positively associated with the high-FN1–CD44 responder state are shown in blue, whereas those negatively associated with FN1–CD44 signaling are shown in red. P values are indicated for significant correlations. (B) Inverse correlation between NFKB1/NF-κB activity and CellChat-derived FN1–CD44 signaling across the six patient–drug conditions (Spearman R = −0.886, two-tailed exact P = 0.0333). Each dot represents one patient–drug condition. The solid line indicates the regression fit, and the shaded region indicates the 95% confidence interval. **(C)** Representative immunofluorescence images showing the spatial association of fibronectin, CD44, and SFRP1 in the synovium-on-a-chip. Scale bars, 50 μm. **(D)** Inverse correlation between the image-derived on-chip FN×CD44 interaction signal and IL-6 secretion across untreated and drug-treated chip conditions (Pearson r = −0.6600, two-tailed P = 0.0054). Each dot represents one patient–condition (16 patient–condition from three patients). The solid line indicates the linear regression fit, and the shaded region indicates the 95% confidence interval. **(E)** Proposed mechanism and fibronectin-rescue strategy. Matrix-retained fibronectin engages CD44 on SFRP1⁺ FLS and restrains NF-κB activity and downstream IL-6 production. Supplementation with exogenous fibronectin is therefore proposed to restore FN–CD44 niche signaling and partially improve TNFi responsiveness in TNFi-refractory chips. **(F)** Quantification of the FN×CD44 interaction signal in chips from patient RA121 treated with TNFi alone or TNFi combined with fibronectin (TNFi + FN) (P = 0.0117). **(G)** IL-6 secretion following TNFi or TNFi + FN treatment on Day 12 (P =0.0258). **(H)** Quantification of collagen IV intensity following TNFi or TNFi + FN treatment (P = 0.0152). **(I)** Integrated on-chip treatment-response score for RA121 following TNFi or TNFi + FN treatment, showing a shift from the predicted no-response category with TNFi alone to the predicted good-response category following fibronectin supplementation. For **(F–H)**, n = 3 chips per condition from one patient; two-tailed unpaired t-test; each dot represents one chip; Bars and error bars indicate mean ± SD.

Among all inferred regulators, nuclear factor kappa-B (*NFKB1*, which encodes NF-κB) showed the strongest negative association, and Nuclear Receptor Subfamily 1 Group H Member 3 (*NR1H3*, which encodes LXRα) the strongest positive association, identifying these two factors as the extremes of the FN1-CD44-associated transcriptional axis rather than as individually selected candidates. *NR1H3* activity increased with FN1–CD44 signaling (Spearman R = 0.943, P = 0.017), whereas *NFKB1* activity decreased (Spearman R = −0.886, P = 0.033; **Fig. 7A, B; Supplemental Fig. 8A**). *NR1H3* and *NFKB1* activities were themselves inversely associated across the same conditions (R = −0.943, P = 0.017; **Supplemental Fig. 8B**). At the gene level, conditions with low FN1-CD44 signaling showed a coordinated upward trend in NF-κB and interferon-associated transcripts such as *NFKBIA* and *IFITM3*, whereas the ECM-protective transcripts *SERPINE2* and *TIMP3* trended in the opposite direction, in line with the inferred TF activities **(Supplemental Fig. 8C)**. These transcriptional patterns further linked loss of ECM-niche signaling with inflammatory activation and motivated direct testing of whether restoring this could improve therapeutic response.

We therefore proposed a model in which matrix-retained FN1 engages CD44 on SFRP1⁺ FLS and maintains an NF-κB-low state possibly from the increased LXRα activity that restrains inflammatory signaling and favors treatment responsiveness. Conversely, loss of FN1–CD44 signaling promotes NF-κB activation and an inflammatory, treatment-resistant stromal phenotype **(Fig. 7E)**. To spatially validate FN–CD44 interactions within the chip, we performed immunofluorescence staining for SFRP1, CD44, and FN **(Fig. 7C).** A 3.2 um (10 pixels)-wide region surrounding the CD44⁺ cell membrane was defined as the perimembrane interaction zone, and the pixel-wise product of CD44 and FN fluorescence within this region was used to calculate an FN×CD44 interaction score (**Supplemental Fig. 8D**). Across three patients evaluated under untreated, IL-6Ra, TNFi, and JAKi conditions, the FN×CD44 interaction score was inversely correlated with IL-6 secretion (Pearson r = −0.6600, P = 0.0054; **Fig. 7D**), supporting a relationship between local FN–CD44 engagement and suppression of inflammatory output.

To test this model directly, we used chips from an independent patient, RA121, which had limited response to TNFi treatment. Exogenous FN was administered together with TNFi for four days and compared with TNFi treatment alone. FN supplementation significantly increased the FN×CD44 interaction signal (P = 0.0117; **Fig. 7F**), reduced secretion of IL-6, an NF-κB-associated inflammatory output (P = 0.0258; **Fig. 7G**), and increased collagen IV intensity, consistent with improved ECM preservation (P = 0.0152; **Fig. 7H**). Consistent with these improvements across individual readouts, the integrated on-chip treatment response score shifted the predicted outcome from no response with TNFi alone to good response following TNFi + FN treatment (**Fig. 7I**). Together, these findings indicate a partial restoration of TNFi responsiveness and support restoration or maintenance of the FN–CD44 ECM niche as a potential strategy for overcoming TNFi treatment resistance.

## Discussion

Chronic inflammatory diseases such as RA are characterized by high inter-patient heterogeneity in both disease pathology and therapeutic response, yet clinicians currently lack reliable guidance on selection among the growing array of treatments. The current trial-and-error treatment strategies are insufficient in the vast majority of patients, who achieve neither remission nor very low disease activity within three to six months of diagnosis. Ineffective treatment results in increased morbidity and unnecessary risk and cost of exposing patients to medications to which they will not respond. Our patient-derived synovial organ-on-a-chip platform could fill this gap by enabling functional drug testing and screening. Despite the pressing clinical need, application of patient-derived organ-on-a-chip technology to drug screening in rheumatic disease remains extremely limited. To our knowledge, ours is the first patient-derived synovium-on-a-chip platform to achieve clinically validated, drug therapeutic prediction across a cohort of RA patients spanning distinct inflammatory endotypes.

By profiling RA patients across defined synovial endotypes, we identified distinct stromal-immune interaction patterns. In high inflammatory endotype chips, sublining FLS accumulated in the perivascular region with a high density of CD90 expression and abundant ECM deposition. Vascular permeability was increased, facilitating immune cell extravasation into the tissue compartment, where immune cells adopted a pro-inflammatory yet exhausted phenotype with limited proliferation. This is similar to findings reported in tumor microenvironments and consistent with other reports of immune aging and cell exhaustion in RA (*40–42*). In contrast, low inflammatory endotype chips were characterized by preserved vascular barrier integrity, where endothelial cells provided survival signals to immune cells that remained predominantly intravascular. These immune cells were actively proliferating but displayed a low-inflammatory phenotype. The highly inflammatory synovium is marked by immune cell recruitment and retention coupled with limited survival, resulting in a dynamic cycle of infiltration, activation, exhaustion, and turnover. These findings establish that the coordinated interplay among vascular cells, FLS, and immune cells collectively determines patient-specific endotype identity, rather than any single cellular compartment acting in isolation.

Multimodal on-chip drug profiling with infliximab, tocilizumab and tofacitinib generated a composite treatment response score integrating four key RA readouts of ECM remodeling, inflammation, immune cell survival, and immune cell recruitment, which correlated strongly with ΔDAS28 (R = 0.844, p=0.0348). Drug response heterogeneity persisted even within the same synovial endotype: among three low inflammatory endotype patients, RA97 responded well to TNFi but not to JAKi, RA99 moderately to TNFi and well to JAKi, and RA105 moderately to JAKi and not to TNFi. Single-cell transcriptomic profiling of these patients across two treatments and untreated controls identified a SFRP1⁺ FLS subpopulation that survived targeted therapy through non-proliferative inflammatory signaling programs. Notably, this drug-resistant population shares key ECM-associated transcriptional features with the COMP⁺ fibrogenic fibroblast subset recently identified in treatment-refractory RA, where expanded fibrogenic niches marked by excessive ECM deposition were linked to non-remission and sustained disease activity despite immune depletion (*43*). CellChat analysis identified FN1-CD44 signaling to SFRP1⁺ FLS as a central axis associated with drug response, with higher connectivity correlating with better treatment response scores. Strong FN1-CD44 engagement directed FLS toward a quiescent, remodeling state, whereas reduced connectivity was associated with sustained inflammatory signaling and drug resistance. Together with the fibrogenic resistance mechanism described by Kartik *et al*.(*43*), these findings position ECM not merely as a structural scaffold but as an active signaling mediator that shapes therapeutic outcome in RA.

Our findings offer a direction that reframes treatment non-response in RA as a form of environment-mediated drug resistance (EMDR). In oncology, EMDR describes a transient, *de novo* resistance phenotype in which tumor cells are protected from therapy through adhesion to extracellular matrix components (cell adhesion-mediated drug resistance, CAM-DR) or paracrine signaling from the stromal microenvironment (*44*). Despite the opposing immune contexts of an immunosuppressive tumor microenvironment and the immune-activated synovium in RA, our findings reveal an analogous but directionally inverted niche mechanism. Whereas ECM adhesion in cancer promotes resistance by protecting malignant cells from elimination, intact ECM signaling in RA appears to constrain pathogenic stromal activation and maintain drug sensitivity. Treatment non-response therefore reflects not simply a failure to eliminate pathogenic cells, but a disruption of ECM-mediated communication among stromal populations. Specifically, intact FN1–CD44 niche signaling maintains SFRP1⁺ FLS in a relatively quiescent state compatible with tissue homeostasis and therapeutic responsiveness. When these niche signals are disrupted under drug pressure, SFRP1⁺ FLS transition toward an inflammatory transcriptional program and expand, establishing a self-reinforcing resistance niche that perpetuates synovial inflammation despite continued therapy. Here, FN1-CD44 interactions do not merely protect cells from death but actively reprogram stromal identity to sustain a low-inflammatory milieu. These findings identify FN1–CD44-mediated stromal niche integrity as a determinant of treatment outcome and motivate strategies aimed at restoring or preserving homeostatic ECM cues, including fibronectin supplementation or prevention of excessive ECM degradation, as potential adjunctive approaches for overcoming EMDR in refractory RA.

The current cohort is sufficient for proof-of-concept demonstration of platform fidelity and treatment response prediction but limits statistical power for establishing generalizable predictive biomarkers. Additionally, the synovial tissue was obtained from patients at the time of knee joint replacement surgery, which introduced selection bias toward advanced disease phenotypes. While autologous blood immune cells provided a scalable source of patient-matched immune cells, tissue-resident immune populations may confer distinct functional properties not fully captured by circulating cells. However, Elmesmari *et al.* recently demonstrated that joint-specific tissue-resident synovial tissue macrophage subsets, including both health-and disease-associated clusters, can derive from peripheral blood monocytes (*45*), which lends support to our strategy. Similarly, endothelial cells in the current platform are HUVEC-derived rather than patient-matched, a common approach in organ-on-chip systems due to the difficulty of isolating sufficient numbers of viable endothelial cells from tissue. Nevertheless, HUVECs co-cultured with endotype-specific FLS and immune cells adopted distinct transcriptional programs between high and low endotypes, suggesting that the patient-specific stromal and immune niche is sufficient to instruct endothelial phenotype in this system. Despite using tissue obtained at the time of total joint arthroplasty and matched peripheral blood immune cells to derive the chips, the multimodal on-chip TNFi and JAKi treatment response data tightly aligned with each patient’s real-world TNFi and JAKi treatment response which typically preceded the tissue sampling by several months or years. This suggests that an individual’s RA biology is imprinted and retained in key aspects captured on chip, even at advanced stages of disease and when therapies have already been given and lends additional support to the imprinting on FLS and circulating immune cells identified in RA (*46, 47*). This also suggests that studying chips derived from patients at any time of disease (e.g., pre-treatment or after failure of initial DMARD treatment) may be helpful.

Our analysis focused primarily on high and low inflammatory endotypes, as these have either tended to be more aggressive, needing intensive therapy (high inflammatory), or have been more resistant to treatment (low inflammatory) (*19, 20, 23, 29, 48–50*). Furthermore, among the tissue obtained at arthroplasty, the mixed endotype tissues exhibited greater heterogeneity in stromal-immune organization, warranting further investigation with expanded cohorts. This is consistent with previous work showing that the histology scoring algorithm derived by machine learning trained by RNAseq data and semi-quantitative scoring of several histologic features was less precise in separating out mixed inflammatory synovial endotype from either high or low inflammatory endotypes than in separating the high or the low endotypes from others (*29*). Future on-chip studies will include characterizing endotype biology in a larger cohort of patients and further study of differentiating treatment response biology among patients. We also plan to study chips derived from RA patients earlier in the disease process, when disease is active and before the need for arthroplasty, and to prospectively test the chip’s predictive capacity for drug selection. Our vascularized patient-derived synovium-on-a-chip platform overcomes limitations of current RA models by recapitulating human synovium and RA disease, retaining patient heterogeneity, scalability (enabling testing of several treatments on serial chips derived from the same patient) and capture of functional readouts that align with real-world treatment response. Major treatment response outputs can be summarized in a composite score and compared among other treatments tested. Thus far, analysis of the data from the RA synovium-on-a-chip has established an ECM-defined stromal niche that shapes immune cell recruitment and activation and identifies environment-mediated drug resistance of FN1-CD44 signaling to SFRP1⁺ FLS. As illustrated by our work, treatment refractory biology can be investigated and potential new treatment strategies revealed. In summary, our vascularized synovium-on-a-chip provides a functional precision medicine framework for RA that bridges population-level molecular profiling and individualized therapeutic decision-making, offering a path toward rational, personalized selection of targeted therapies.

## MATERIALS AND METHODS

### Study design

This study aimed to develop a physiologically relevant, patient-derived, immunocompetent human synovium-on-a-chip platform to reconstruct key structural and functional features of the RA synovial microenvironment, including lining–sublining spatial organization, vascularized stromal architecture, extracellular matrix remodeling, inflammatory signaling, and immune cell recruitment. Using FLS isolated from RA patient synovial tissues and matched peripheral blood immune cells, we established an individualized and vascularized RA synovium-on-a-chip system to model patient-specific stromal–immune interactions and therapeutic responses.

RA patients were clinically assessed and stratified into low, mixed, and high inflammatory synovial endotypes using a machine-learning algorithm that incorporates the Hospital for Special Surgery synovitis score and gene expression data as previously published by Orange et al. (*29*). We first evaluated whether the chip platform recapitulated RA-specific synovial architecture and inflammatory features by integrating immunofluorescence imaging, cytokine profiling, extracellular matrix analysis, immune cell phenotyping, and single-cell RNA sequencing. We then investigated how endotype-specific stromal–vascular programs regulate immune cell localization, survival, activation, and recruitment within the synovial niche. To assess the translational potential of the platform for personalized therapeutic response prediction, patient-derived chips were treated with clinically relevant RA therapeutics, including infliximab, tocilizumab, and tofacitinib. Drug-induced changes in inflammatory cytokine production, ECM remodeling, immune survival, and immune recruitment were integrated into a composite on-chip treatment response score and compared with available clinical treatment response data.

To further dissect mechanisms of therapeutic response and resistance, we performed single-cell transcriptomic profiling of drug-treated and untreated chips from three low-inflammatory endotype patients with divergent on-chip responses to TNF and JAK inhibition. This analysis was used to identify drug-resistant FLS states and cell–cell communication pathways associated with treatment outcome. All chip experiments were performed using independent biological patient samples and multiple technical chip replicates as sample availability allowed. Experiments were not blinded unless otherwise stated. No data were excluded unless there was an obvious technical failure, including chip leakage, failed gel loading, bacterial contamination, imaging failure, or insufficient cell recovery for downstream analysis. No statistical methods were used to predetermine sample size. All procedures involving human participants were approved by the appropriate Institutional Review Board and conducted in accordance with the principles of the Declaration of Helsinki. Written informed consent was obtained from all participants. All patient samples were deidentified before analysis to ensure compliance with institutional and HIPAA regulations.

### Human cohort

Human synovial tissue and matched peripheral blood samples were obtained from patients with RA undergoing standard of care total knee arthroplasty. Samples were obtained from osteoarthritis patients at the time of standard of care total knee arthroplasty as non-inflammatory disease controls. RA diagnosis was established by the treating rheumatologist and verified by a rheumatologist on our study team by chart review and patient interview, using the Rheumatology (ACR)/European League Against Rheumatism 2010 classification criteria for RA (*51*) and the ACR 1987 criteria (*52*). Clinical information including tender joint count, swollen joint count, erythrocyte sedimentation rate (ESR), C-reactive protein (CRP), patient global health assessment, and clinical treatment history were obtained from detailed review of medical records when available and accessible. DAS28-ESR and EULAR treatment response (*39*) over time were calculated from the data.

Nine RA patients spanning low, mixed, and high inflammatory endotypes were included for platform characterization, inflammatory endotype analysis, and on-chip drug profiling with infliximab, tocilizumab, and tofacitinib. Five RA patients had real-world treatment response data readily available; five had been treated with TNFi and one had been treated with JAKi. Three of these patients shared the same endotype (low-inflammatory) yet had divergent drug response profiles. These were selected for single-cell RNA sequencing of untreated and drug-treated chips to investigate mechanisms of therapeutic resistance. An additional RA patient was included for functional evaluation of treatment resistant niche modulation. All samples were deidentified before analysis. Age of RA patient range was 55-77 (average 65) years. Ninety percent of patients were positive for either rheumatoid factor and/or anti-citrullinated peptide antibody. Ninety percent were female. Seventy percent were Caucasian, twenty percent Black, ten percent Asian. Twenty percent were Hispanic or Latino. Available clinical information and grouping by endotype is summarized in **Supplemental Table 1**.

### Synovial tissue and PBMC isolation

Synovial tissue was obtained from the surgeon during total knee arthroplasty and then examined by an expert musculoskeletal pathologist. Specimen suitable for downstream analysis was identified, transferred to our laboratory and immediately processed. Tissue was cut into fragments and cryopreserved until use. On the date of surgery, matched peripheral blood mononuclear cells (PBMCs) were isolated from whole blood and cryopreserved until use.

### Cell culture

Human Umbilical Vein Endothelial Cells (HUVECs; Lonza) were cultured in EGM-2 medium (Lonza). Patient-derived FLS were isolated from synovial tissue by mechanical and enzymatic dissociation (Supplemental Methods) and maintained in Synoviocyte Growth Medium (Sigma-Aldrich). All primary FLS and HUVECs were used between passages 2 and 4.

Before chip loading, PBMCs were thawed and CD3⁺ T cells, CD19⁺ B cells, and monocytes were isolated by magnetic-activated cell sorting and monocyte-derived macrophages were prepared as described in the **Supplemental Methods**. Cell loading densities were standardized across experiments (**Supplemental Methods**).

### Device design, fabrication, and cell loading

The microfluidic devices were fabricated using standard soft lithography replica molding of PDMS (SYLGARD 184, Dow Corning) from a photolithographically prepared SU-8 master mold (**Supplemental Fig. 1B**). Detailed fabrication and hydrogel preparation protocols are provided in the Supplemental Methods. The 3D microfluidic-based RA synovium-on-a-chip device is composed of three distinct functional regions: a central synovial fluid region, an inner ring representing the lining region, and an outer ring representing the sublining region. The device incorporates fibrin hydrogel to maintain 3D cell culture. Additionally, two side ports are included for optional immune cell loading to simulate immune cell recruitment. Four medium reservoirs are equipped to ensure a consistent culture medium supply and waste removal.

After the PDMS microfluidic devices was made, RAFLSs and macrophages were embedded in the lining region, whereas T cells, B cells, monocytes, and endothelial cells were incorporated into the sublining region within a 3D biomimetic fibrin hydrogel (3 mg/mL for inner ring; 4.5 mg/mL for outer ring). A multistep loading protocol was used to spatially encapsulate RAFLSs and monocyte-derived macrophages within the lining compartment and seed T cells, B cells, monocytes, and ECs in the sublining compartment. Chips were cultured for 8 days to allow tissue maturation, followed by an additional 4 days under treatment conditions for response assessment.

### Immunofluorescence staining

Chips were fixed with fixation buffer (BioLegend) at defined time points and stained with fluorophore-conjugated antibodies targeting FLS markers (CD90, PRG4, PDPN, FAP), vascular cells (CD31), hematopoietic cells (CD45), T cells (CD3, CD4, CD8, RORyt, T-bet), B cells (CD19, CD27, CD38), and monocytes/macrophages (CD14, CD16, HLA-DR, CD163, CD68). For intracellular targets (Ki67), chips were permeabilized with 0.3% Triton X-100 (Sigma-Aldrich) for 30 minutes prior to staining. ECM deposition was assessed by Alexa Fluor 647-Collagen IV (Thermo Fisher Scientific) staining. All chips were counterstained with DAPI (Thermo Fisher Scientific) and imaged using a Nikon C2i confocal microscope. Mean fluorescence intensity was quantified using Qupath or Fiji (version 2). Immune cell subtype classification based on fluorescence intensity gating is described in the Supplemental Methods. A complete list of antibodies, catalogue numbers, and working dilutions is provided in **Supplemental Table 2**.

### Cytokine profiling and quantification

Supernatants were collected from RA chips at defined time points, centrifuged at 2,000 × g for 10 min at 4 °C to remove cellular debris, and stored at −20 °C. Qualitative cytokine profiles were assessed using a Human Cytokine Array C5 membrane kit (catalogue AAH-CYT-5-8, RayBiotech) according to the manufacturer’s protocol. Quantitative measurement of IL-6 concentration was performed using a Human IL-6 ELISA MAX Deluxe kit (catalogue number 430515, BioLegend). MMP secretion was quantified using a Human MMP Array C1 (catalogueAAH-MMP-1-8, RayBiotech). Chemiluminescent signals were captured on a ChemiDoc imaging system (Bio-Rad) and spot intensities were quantified in ImageJ using the Protein Array Analyzer plug-in.

### On-chip drug treatment and response assessment

After 8 days of culture, RA synovium-on-a-chips were treated with infliximab (TNF inhibitor, 10 µg/ml; catalogue HY-P9970, MCE), tocilizumab (IL-6 receptor antagonist, 10 µg/ml; catalogue HY-P9917, MCE), or tofacitinib (JAK inhibitor, 10 µg/ml; catalogue HY-40354, MCE) for 4 days. Untreated chips from the same patient served as matched controls. Media (with treatment for treated chips or without treatment for untreated chips) was refreshed daily. Morphological changes were evaluated by immunofluorescence staining on days 2, 4, 8, 10, and 12. Culture supernatants were collected daily for downstream immunoassays. Drug-induced changes (Δ values) were calculated by subtracting the untreated readout from the treated readout for each of four multimodal readout channels: immune cell migration distance (recruitment), IL-6 concentration (inflammation), CD45⁺ cell count (immune survival), and Collagen IV intensity (ECM remodeling). Each of the four readouts was z-score–normalized across the three treatments within each patient; the composite treatment response score was then computed as the mean of the four z-scored readouts. The detailed mathematical procedure for calculating the score is described in the **Supplementary Methods**. Applying established EULAR treatment response criteria (good response: ΔDAS28 > 1.2; no response: ΔDAS28 < 0.6) (*39*) to a linear prediction model derived from clinical validation (Y = 3.444x + 0.505), each patient’s on-chip response to each drug was classified as good, moderate, or no response.

### Fibronectin supplementation

Human plasma fibronectin (Corning, CLS354008) was added to the perfusion medium at 40 μg/mL together with TNFi (Infliximab,10 μg/ml) for four days and replenished daily. Fibronectin was administered in soluble form rather than as a pre-adsorbed coating. Control chips received TNFi and an equivalent volume of PBS. The experiment was performed using FLS and PBMCs from an independent patient, RA121, who had mixed inflammatory synovial endotype and limited on-chip response to TNFi. TNFi and TNFi+fibronectin chips were generated from the patient and processed in parallel.

### Single-cell RNA sequencing

To compare on-chip and tissue profiles, synovial tissue from patients RA98 and RA105 and cells retrieved on day 8 from synovium-on-a-chips derived from the same patients were processed for scRNA-seq (dataset 1). To assess patient-specific drug responses, chips derived from three patients (RA97, RA99, and RA105) under untreated, TNFi-treated, and JAKi-treated conditions on Day 12 were processed for scRNA-seq (dataset 2). Single-cell suspensions were prepared by enzymatic dissociation, labelled with TotalSeq anti-human hashtag antibodies (BioLegend; 1:250 dilution), and pooled for multiplexed capture, with one pooled library per dataset. Libraries were prepared using the Chromium Single Cell 3′ Reagent Kit v3 (10x Genomics; Single Cell 3′ Library & Gel Bead Kit v3, PN-1000075; Chip Kit v3, PN-1000073; i7 Multiplex Kit, PN-120262) following the manufacturer’s instructions (CG000183 Rev B). Sequencing was performed on an Illumina NovaSeq 6000 with 28-bp read 1, an 8-bp i7 index, and 91-bp read 2.

Raw sequencing data were processed using Cell Ranger v7.0 (10x Genomics) and aligned to the GRCh38 human reference genome. Both datasets were analyzed using a common workflow in Seurat v5. Hashtag counts were normalized and demultiplexed, and only singlets were retained. Genes detected in fewer than three cells were excluded. For dataset 1, cells were retained with 200–6,200 detected genes and fewer than 28,000 UMIs for chip samples, or 200–5,800 detected genes and fewer than 22,000 UMIs for tissue samples, with less than 18% mitochondrial reads in both sample types. For dataset 2, cells were retained with 200–6,500 detected genes, fewer than 30,000 UMIs, and less than 10% mitochondrial reads. Expression data were normalized using SCTransform v2 with 3,000 variable features. The first 20 principal components were used for UMAP visualization and shared nearest neighbour graph-based Louvain clustering, with resolutions of 0.7 and 0.5 for datasets 1 and 2, respectively. Clusters were annotated using canonical markers. In dataset 1, five clusters absent from chip samples were excluded from subsequent comparisons, leaving 22,942 cells. Dataset 2 contained 42,335 cells and included eight FLS clusters annotated as distinct functional states. FLS were identified based on expression of known markers and subset for downstream analysis. Eight FLS functional states were annotated on the basis of differentially expressed genes and published marker gene sets: IFN-activated (*ISG15*, *IFIT1*, *MX1*), Structural-remodeling (*SFRP1*, *SOX9*, *SCARA5*), ECM-IL1 Remodeling, Lipid metabolic, Recruitment lining, IL-6⁺ recruitment, Proliferative, and Basal FLS. Differential gene expression between conditions was performed using the Wilcoxon rank-sum test with Bonferroni correction for genes detected in at least 10% of cells (adjusted P < 0.05, |log2FC| > 0.25). Sex-linked genes (Y-chromosome genes and XIST/RPS4X) were excluded from differential expression and gene set enrichment analyses to avoid confounding by patient sex.

### Statistics and reproducibility

Details regarding biological and technical replicates, as well as repetitions for each experiment, are listed in the respective figure legends. Data are presented as mean ± standard deviation (SD), unless otherwise specified. Representative images were selected from the independent samples or chips specified in the corresponding figure legends. Statistical analyses were performed using GraphPad Prism v10 and R v4.3. Comparisons between two groups used two-tailed unpaired Student’s t-tests, Welch’s t-tests, or Wilcoxon rank-sum tests, as specified in the figure legends. Where indicated, data were log10-transformed before analysis. Comparisons among multiple groups used one-way or two-way analysis of variance (ANOVA), followed by Dunnett’s or Šídák’s multiple-comparisons tests, as appropriate. Statistical procedures for scRNA-seq analyses are described in the relevant Methods sections and figure legends; Benjamini– Hochberg correction was applied where indicated. Associations were assessed using Pearson or Spearman correlation with two-tailed P value and are described in the relevant figure legends. Leave-one-out cross-validation was used to evaluate the robustness of the linear prediction model relating on-chip composite response scores to clinical ΔDAS28 values. Adjusted P values were computed using the Benjamini–Hochberg method unless otherwise stated. No statistical methods were used to predetermine sample size. Typically, receipt of records to calculate real-world Eular response lagged behind drug testing on chip. The investigators were not blinded to patient endotype during chip fabrication and drug treatment.

## SUPPLEMENTAL MATERIALS

Supplemental Methods

Supplemental Figs. 1 to 8

Supplemental Tables 1 to 2

## ACKNOWLEDGEMENTS

We thank all patients who generously donated synovial tissue and peripheral blood samples for this study. We thank the Hospital for Special Surgery surgical teams, research assistants (with special thanks to Sandy Zeng, Azka Ali, and Sanah Chatiwala), and rheumatologists for their assistance with patient recruitment, sample collection, and clinical data coordination. We thank Drs. Romy Kallas, Amit Lakhanpal, and Melanie Smith for assistance with determination of tender and swollen joint counts on the date of arthroplasty. We thank Drs. Edward DiCarlo, Daniel Ramirez, and Yaxia Zhang for assistance with synovial tissue selection from surgical specimens for downstream processing; Drs. Janet Sun and Dena Almeida for support with tissue microscopy imaging; and Dr. Chendong Peng and Weill Cornell Medicine Genomics Core Facility for assistance with single-cell RNA sequencing library preparation and sequencing

## Author contributions

T.W.M., R.C., T.N., and W.C. conceived the project. R.C., T.W.M., T.N., and W.C. designed the experiments. T.W.M. verified patient diagnoses, obtained and characterized patient synovial tissue and peripheral blood samples, retrieved clinical data, and calculated EULAR treatment responses. Y.L. and C.F. assisted with patient identification, recruitment, and retrieval of clinical records; Y.L. also assisted with record review and calculation of DAS-28; C.H. assisted with sample processing and quality control. S.R. provided first synovial tissue samples from joint surgery which allowed for key synovium-on-a-chip advancements; D.M. provided assistance in obtaining samples during standard of care TKAs; D.R. provided expert histopathological scoring of the synovial tissues. K.K. provided additional engineering expert review and appraisal of this work. R.C. led the on-chip experimental work. F.S. and L.L. assisted with fabrication of the chip molds. E.S. and E.W. assisted with selected on-chip experiments. R.C., T.W.M., T.N., and W.C. contributed to data interpretation. The manuscript was drafted by R.C. and T.W.M. and was reviewed and edited by W.C. and T.N. All authors reviewed and accepted the final version of this manuscript

## Data and materials availability

All data associated with this study are present in the paper or the Supplemental Materials. Single-cell RNA sequencing raw data reported in this paper will be deposited in the Gene Expression Omnibus. The human GRCh38 reference genome used for scRNA-seq read alignment is available. Additional processed data, analysis scripts, and materials are available from the corresponding author upon reasonable request

## References

1. G. S. Firestein, I. B. McInnes, Immunopathogenesis of Rheumatoid Arthritis. Immunity 46, 183–196 (2017).

2. F. Zhang et al., Defining inflammatory cell states in rheumatoid arthritis joint synovial tissues by integrating single-cell transcriptomics and mass cytometry. Nat Immunol 20, 928-+ (2019).

3. S. E. Gabriel, The epidemiology of rheumatoid arthritis. Rheum Dis Clin N Am 27, 269–281 (2001).

4. J. M. Gwinnutt et al., Twenty-Year Outcome and Association Between Early Treatment and Mortality and Disability in an Inception Cohort of Patients With Rheumatoid Arthritis Results From the Norfolk Arthritis Register. Arthritis & Rheumatology 69, 1566–1575 (2017).

5. T. Sokka et al., Work disability remains a major problem in rheumatoid arthritis in the 2000s: data from 32 countries in the QUEST-RA study. Arthritis Res Ther 12, R42 (2010).

6. M. Boers, Understanding the window of opportunity concept in early rheumatoid arthritis. Arthritis Rheum 48, 1771–1774 (2003).

7. J. S. Smolen et al., Treating rheumatoid arthritis to target: 2014 update of the recommendations of an international task force. Annals of the Rheumatic Diseases 75, 3–15 (2016).

8. J. S. Smolen et al., Rheumatoid arthritis. Nat Rev Dis Primers 4, 18001 (2018).

9. L. Ouboussad, A. N. Burska, A. Melville, M. H. Buch, Synovial Tissue Heterogeneity in Rheumatoid Arthritis and Changes With Biologic and Targeted Synthetic Therapies to Inform Stratified Therapy. Front Med (Lausanne) 6, 45 (2019).

10. A. H. V. V. Mil, Synovial tissue for personalised medicine in rheumatoid arthritis: hype or hope? Lancet Rheumatol 5, e640–e641 (2023).

11. J. J. Liu et al., Ivarmacitinib, a selective Janus kinase 1 inhibitor, in patients with moderate-to-severe active rheumatoid arthritis and inadequate response to conventional synthetic DMARDs: results from a phase III randomised clinical trial. Annals of the Rheumatic Diseases 84, 188–200 (2025).

12. C. J. van de Laar et al., PERFECTRA: a pragmatic, multicentre, real-life study comparing treat-to-target strategies with baricitinib versus TNF inhibitors in patients with active rheumatoid arthritis after failure on csDMARDs. Rmd Open 10, (2024).

13. A. Floudas et al., Distinct stromal and immune cell interactions shape the pathogenesis of rheumatoid and psoriatic arthritis. Annals of the Rheumatic Diseases 81, 1224–1242 (2022).

14. A. P. Croft et al., Distinct fibroblast subsets drive inflammation and damage in arthritis. Nature 570, 246–251 (2019).

15. H. Yang et al., Single-Cell Profiling Identifies Il1bhi Macrophages Associated with Inflammation in Pd-1 Inhibitor-Induced Inflammatory Arthritis. Annals of the Rheumatic Diseases 83, (2024).

16. X. Y. Xia et al., Single cell immunoprofile of synovial fluid in rheumatoid arthritis with TNF/JAK inhibitor treatment. Nat Commun 16, (2025).

17. F. Zhang et al., Deconstruction of rheumatoid arthritis synovium defines inflammatory subtypes. Nature 623, 616-+ (2023).

18. F. Humby et al., Rituximab versus tocilizumab in anti-TNF inadequate responder patients with rheumatoid arthritis (R4RA): 16-week outcomes of a stratified, biopsy-driven, multicentre, open-label, phase 4 randomised controlled trial. Lancet 397, 305–317 (2021).

19. M. J. Lewis et al., Molecular Portraits of Early Rheumatoid Arthritis Identify Clinical and Treatment Response Phenotypes. Cell Reports 28, 2455-+ (2019).

20. M. H. Buch, A. Melville, D. G. McGonagle, Synovial cellular and molecular signatures stratify clinical response to csDMARD therapy and predict radiographic progression in early rheumatoid arthritis patients. Annals of the Rheumatic Diseases 79, (2020).

21. F. Rivellese et al., Rituximab versus tocilizumab in rheumatoid arthritis: synovial biopsy-based biomarker analysis of the phase 4 R4RA randomized trial. Nat Med 28, 1256-+ (2022).

22. M. Rehberg et al., Identification of a Rule to Predict Response to Sarilumab in Patients with Rheumatoid Arthritis Using Machine Learning and Clinical Trial Data (Sep, 10.1007/s40744-021-00361-5, 2021). Rheumatol Ther 8, 1921–1922 (2021).

23. G. Dennis, Jr., et al., Synovial phenotypes in rheumatoid arthritis correlate with response to biologic therapeutics. Arthritis Res Ther 16, R90 (2014).

24. D. E. Ingber, Challenges and opportunities for human Organ Chips in FDA assessments and pharma pipelines. Cell Stem Cell 33, 176–183 (2026).

25. E. M. L. Philippon et al., A novel 3D spheroid model of rheumatoid arthritis synovial tissue incorporating fibroblasts, endothelial cells, and macrophages. Frontiers in Immunology 14, (2023).

26. M. Rothbauer et al., Establishment of a human three-dimensional chip-based chondro-synovial coculture joint model for reciprocal cross talk studies in arthritis research. Lab on a Chip 21, 4128–4143 (2021).

27. W. Diao et al., Synovium-On-A-Chip: Simulating the Microenvironment of the Rheumatoid Arthritis Synovium via Multicell Interactions to Target Fibroblast-Like Synoviocytes. Adv Sci (Weinh) 12, e11945 (2025).

28. C. L. Thompson et al., Human vascularised synovium-on-a-chip: a mechanically stimulated, microfluidic model to investigate synovial inflammation and monocyte recruitment. Biomed Mater 18, (2023).

29. D. E. Orange et al., Identification of Three Rheumatoid Arthritis Disease Subtypes by Machine Learning Integration of Synovial Histologic Features and RNA Sequencing Data. Arthritis & Rheumatology 70, 690–701 (2018).

30. J. S. Smolen, D. Aletaha, I. B. McInnes, Rheumatoid arthritis. Lancet 388, 2023–2038 (2016).

31. B. Bartok, G. S. Firestein, Fibroblast-like synoviocytes: key effector cells in rheumatoid arthritis. Immunol Rev 233, 233–255 (2010).

32. I. Korsunsky et al., Cross-tissue, single-cell stromal atlas identifies shared pathological fibroblast phenotypes in four chronic inflammatory diseases. Med-Cambridge 3, 481-+ (2022).

33. D. A. Rao et al., Pathologically expanded peripheral T helper cell subset drives B cells in rheumatoid arthritis. Nature 542, 110-+ (2017).

34. S. Alivernini et al., Distinct synovial tissue macrophage subsets regulate inflammation and remission in rheumatoid arthritis. Nat Med 26, 1295–1306 (2020).

35. I. B. McInnes, G. Schett, The pathogenesis of rheumatoid arthritis. N Engl J Med 365, 2205–2219 (2011).

36. U. Hardt et al., Integrated single cell and spatial transcriptomics reveal autoreactive differentiated B cells in joints of early rheumatoid arthritis. Sci Rep 12, 11876 (2022).

37. K. V. Wei et al., Notch signalling drives synovial fibroblast identity and arthritis pathology. Nature 582, 259-+ (2020).

38. S. Turcinov et al., Diversity and Clonality of T Cell Receptor Repertoire and Antigen Specificities in Small Joints of Early Rheumatoid Arthritis. Arthritis & Rheumatology 75, 673–684 (2023).

39. J. Fransen, P. L. van Riel, The Disease Activity Score and the EULAR response criteria. Rheum Dis Clin North Am 35, 745–757, vii-viii (2009).

40. Y. Zhao, Q. Shao, G. Peng, Exhaustion and senescence: two crucial dysfunctional states of T cells in the tumor microenvironment. Cell Mol Immunol 17, 27–35 (2020).

41. A. Chin, A. Small, S. W. Wong, M. D. Wechalekar, T Cell Dysregulation in Rheumatoid Arthritis: from Genetic Susceptibility to Established Disease. Curr Rheumatol Rep 27, 14 (2025).

42. T. Huang et al., Macrophage metabolic exhaustion and PANoptotic cell death drive chronic tissue inflammation in rheumatoid arthritis. Immunity 59, 970–987 e976 (2026).

43. K. Bhamidipati et al., Spatial patterning of fibroblast TGFbeta signaling underlies treatment resistance in rheumatoid arthritis. Nat Immunol 27, 556–571 (2026).

44. M. B. Meads, R. A. Gatenby, W. S. Dalton, Environment-mediated drug resistance: a major contributor to minimal residual disease. Nat Rev Cancer 9, 665–674 (2009).

45. A. Elmesmari et al., The ontogeny of myeloid-stromal synovial tissue niches in rheumatoid arthritis. Res Sq, (2025).

46. G. Nygaard, G. S. Firestein, Restoring synovial homeostasis in rheumatoid arthritis by targeting fibroblast-like synoviocytes. Nat Rev Rheumatol 16, 316–333 (2020).

47. W. Wang et al., Multi-lineage transcriptional and cell communication signatures define pathways in individuals at-risk for developing rheumatoid arthritis that initiate and perpetuate disease. Res Sq, (2025).

48. A. Nerviani et al., A Pauci-Immune Synovial Pathotype Predicts Inadequate Response to TNFalpha-Blockade in Rheumatoid Arthritis Patients. Front Immunol 11, 845 (2020).

49. F. Rivellese et al., Stratification of biological therapies by pathobiology in biologic-naive patients with rheumatoid arthritis (STRAP and STRAP-EU): two parallel, open-label, biopsy-driven, randomised trials. Lancet Rheumatol 5, e648–e659 (2023).

50. F. Humby et al., Synovial cellular and molecular signatures stratify clinical response to csDMARD therapy and predict radiographic progression in early rheumatoid arthritis patients. Ann Rheum Dis 78, 761–772 (2019).

51. D. Aletaha et al., 2010 rheumatoid arthritis classification criteria: an American College of Rheumatology/European League Against Rheumatism collaborative initiative. Ann Rheum Dis 69, 1580–1588 (2010).

52. F. C. Arnett et al., The American Rheumatism Association 1987 revised criteria for the classification of rheumatoid arthritis. Arthritis Rheum 31, 315–324 (1988).

