## Supplemental Methods; Supplemental Figs. 1 to 8; Supplemental Tables 1 to 2 for "Patient-Derived Synovium-on-a-Chip Recapitulates Disease and Treatment Response and Permits Study of Resistance in Rheumatoid Arthritis"

**The PDF file includes:**

Supplementary Methods
Supplementary Figures 1 to 8
Supplementary Tables. 1 to 2

### Supplementary Methods

#### Synovial tissue processing and FLS isolation

Human synovial tissue specimens were obtained from RA and OA patients undergoing clinically indicated total joint arthroplasty at the Hospital for Special Surgery (HSS). All procedures involving human tissues were approved by the Institutional Review Board at HSS and were conducted in accordance with the principles of the Declaration of Helsinki. All participants provided written informed consent. All samples were deidentified before analysis to ensure compliance with HIPAA regulations. Synovial tissue fragments were dissociated mechanically and enzymatically in digestion buffer containing Liberase TL (100 μg/mL; Sigma-Aldrich) and DNase I (100 μg/mL; New England Biolabs) in RPMI on a rotator at 37°C for 30 minutes. The resulting cell suspension was transferred to RPMI supplemented with 10% heat-inactivated fetal bovine serum (FBS; Invitrogen) and 1% glutamine (Thermo Scientific) and cultured for 10 days as passage 0 cells. Primary FLS were identified by adherent spindle-shaped morphology and used for experiments between passages 2 and 4. For histological assessment, frozen synovial tissue sections were prepared at 5 μm thickness and subjected to hematoxylin and eosin staining for histopathological scoring.

#### PBMC isolation and immune cell preparation

Matched PBMCs were isolated from whole blood by density gradient centrifugation and cryopreserved at $5 \times 10^6$ cells per vial in CryoStor CS10 (Sigma-Aldrich) per manufacturer instructions. Three days before chip loading, one vial ($5 \times 10^6$ cells) was thawed for T cell and macrophage preparation. CD3+ T cells were isolated using the EasySep™ Release Human CD3 Positive Selection Kit (StemCell Technologies) and activated with ImmunoCult™ Human CD3/CD28 T Cell Activator according to the manufacturer's instructions. Cells were cultured in RPMI 1640 supplemented with 10% FBS for 3 days and harvested on day 3. Monocytes were isolated from the remaining PBMC fraction using the MojoSort™ Human Pan Monocyte Isolation Kit (BioLegend, 480059) and cultured in RPMI supplemented with 50 ng/mL macrophage colony-stimulating factor (M-CSF; PeproTech) for 3 days to induce macrophage differentiation. Macrophages were collected on day 3. For the OA–RA comparisons shown in Fig. 2, monocytes were differentiated with M-CSF for 5 days, followed by polarization with 20 ng/mL IFN-γ and 10 pg/mL LPS for an additional 3 days. The resulting M1-polarized macrophages were harvested on day 8 for chip loading. On the day of chip loading, an additional vial ($5 \times 10^6$ PBMCs) was thawed. $CD19^+$ B cells were isolated using the EasySep™ Release Human CD19 Positive Selection Kit (StemCell Technologies), and monocytes were subsequently isolated from the remaining PBMC fraction as described above.

#### Hydrogel preparation

A gelatin solution (12mg/mL) was prepared by dissolving gelatin powder derived from porcine skin (G2500, Sigma-Aldrich) in 1× Dulbecco's phosphate-buffered saline (DPBS, without calcium and magnesium;

Invitrogen). The solution was heated and stirred at 37°C for 30 minutes, followed by sterile filtration, aliquoting, and storage at 4°C. The fibrinogen solution (9 mg/mL for outer ring; 6 mg/mL for inner ring) was prepared by dissolving lyophilized fibrinogen from bovine plasma (F8630, Sigma-Aldrich) in DPBS at 37°C for 1 hour. The sterile-filtered fibrinogen solution was stored at 4°C for use within one week. The thrombin solution was prepared by reconstituting lyophilized thrombin (604980, Sigma-Aldrich) in sterile DPBS at a concentration of 100 U/mL and stored in aliquots at −20°C.

**Microfluidic chip fabrication**

The microfluidic device was fabricated using a standard soft lithography replica molding technique. The mold was created using SU-8 negative photoresist (2050, MicroChem) at a thickness of 100 μm on a silicon wafer by photolithography. Before replica molding, the surface of the SU-8 mold was treated overnight in a vacuum desiccator with trichloro(1H,1H,2H,2H-perfluorooctyl)silane (448931, Sigma-Aldrich) vapor to facilitate PDMS release. A mixture of PDMS base and curing agent (SYLGARD 184, Dow Corning) at a 10:1 weight-to-weight ratio was thoroughly mixed, cast onto the mold, degassed, and cured at 80°C for 1 hour. Once cured, the PDMS layer was peeled from the mold, and 1- and 3-mm holes were punched to create the inlets and medium reservoirs, respectively. The PDMS slabs were cleaned and bonded to glass coverslips (22 mm × 22 mm; Thermo Fisher Scientific) using oxygen plasma (PE-50 XL, Plasma Etch; 350 W, 2 minutes), followed by overnight incubation at 80°C to restore hydrophobicity. Assembled devices were sterilized under ultraviolet light in a type 2 class laminar flow hood for 20 minutes before cell loading.

**Cell loading and on-chip culture**

All experiments were performed using serial chips derived from patient biosamples. For each patient, approximately 40 chips were allocated for longitudinal characterization of on-chip features across multiple time points, and an additional 48 chips were designated for drug screening across four conditions (untreated control, tocilizumab, infliximab, and tofacitinib). Each experimental condition was performed with a minimum of three biological replicates (independent chips seeded from the same patient sample), and ten randomly selected fields per chip were imaged as technical replicates for quantitative analysis. RA-FLS and macrophages were embedded in the lining region, whereas endothelial cells were incorporated into the sublining region within a 3D biomimetic fibrin hydrogel (9 mg/mL for outer ring; 6 mg/mL for inner ring). T cells, B cells, and monocytes from matched patient PBMCs were loaded into the sublining region along with the endothelial cells at day 0. Although PBMC composition varied slightly among patients, final cell loading concentrations were standardized across experiments. The approximate cell densities used for chip loading were as follows: HUVECs, $1.5 \times 10^7$ cells/mL; RA-FLS, $2.5 \times 10^6$ cells/mL; macrophages, $5 \times 10^5$ cells/mL; activated $CD3^+$ T cells, $2 \times 10^5$ cells/mL; $CD19^+$ B cells, $1 \times 10^6$ cells/mL; and monocytes, $1 \times 10^6$ cells/mL. Chips were cultured in a mixed medium consisting of EGM-2 (Lonza) and Synoviocyte Growth Medium (Sigma-Aldrich) at a 2:1 ratio. Chips were maintained for 8 days to allow tissue maturation, followed by an additional 4 days under treatment conditions for treatment response assessment. Culture medium was refreshed daily.

**Time-lapse live microscopy**

To visualize immune cell recruitment and migration dynamics on chip, immune cells isolated from one healthy donor's peripheral blood were labeled with CellTracker™ Fluorescent Probes (Thermo Fisher Scientific, V22887) prior to loading into the side channel. Chips were mounted on an inverted microscope equipped with a motorized stage and an environmental control chamber (Okolab; 37°C, 5% $CO_2$). Phase-contrast and fluorescence images were acquired every 5 minutes for 2 hours using a digital CMOS camera (ORCA-Flash4.0 LT, Hamamatsu Photonics) with a 10× objective. Immune cell migration parameters were quantified by tracking individual cells (>75 cells per condition) in ImageJ using the Manual Tracking plug-in. Average migration speed was defined as the total distance traveled divided by observation time.

**Immunofluorescence staining**

Chips were fixed with fixation buffer (BioLegend, 420801) for 30 minutes at room temperature and washed three times with PBS, 10 minutes each. For intracellular staining requiring membrane permeabilization, chips were incubated with 0.3% Triton X-100 (Sigma-Aldrich, 11332481001) in PBS for 30 minutes at room temperature, followed by PBS washes. Primary antibodies were diluted in Cell Staining Buffer (BioLegend, 420201). Chips were incubated with fluorophore-conjugated primary antibodies and DAPI (1 µg/mL; Thermo Fisher Scientific, D1306) overnight at 4°C in the dark. After staining, chips were washed with PBS, kept in Cell Staining Buffer and imaged using a Nikon C2i confocal microscope. Mean fluorescence intensity was quantified using Fiji (version 2) or Qupath. A complete list of antibodies, catalogue numbers, clone information, and working dilutions is provided in Supplementary Table S2.

**Image-based immune cell subtype classification**

Immune cell subtype classification on chip was performed using fluorescence intensity-based gating of confocal immunofluorescence images, analogous to conventional flow cytometry analysis. All chips used for immune phenotyping were fixed, stained, and imaged under same antibody concentrations, laser power, gain, and exposure settings to ensure consistency across conditions **(Supplementary Fig. 7)**.

*T cell subtype classification*. CD4+ T cells were identified by CD4 and CD45 co-expression. T cell subsets were classified based on T-bet and ROR-γt fluorescence intensity. Classification thresholds were determined from the normalized frequency distribution of T-bet and ROR-γt intensity in CD4+ cells relative to unlabeled controls. Cells exceeding the T-bet threshold were classified as Th1 cells, cells exceeding the ROR-γt threshold were classified as Th17 cells, and remaining CD4+ cells were classified as unpolarized.

*B cell subtype classification*. B cells were identified as CD19$^+$ populations. Subsets were classified based on CD27 and CD38 fluorescence intensity relative to CD19$^-$ cells in the same field of view, which served as internal negative controls for background signal estimation. To allow consistent comparison across conditions, fluorescence intensities from CD19$^+$ and CD19$^-$ cells were pooled to establish unified thresholds. Cells were categorized as CD27$^+$ or CD38$^+$ based on thresholds exceeding the pooled mean, and as CD38$^{++}$ if exceeding the mean plus two standard deviations. Naive B cells were defined as CD27$^-$CD38$^-$, memory B cells as CD27$^+$CD38$^{low}$, and plasma cells as CD38$^{high}$. Scatter plots of CD27 versus CD38 were generated for visualization, and subset percentages were quantified using MATLAB R2024b.

*Monocyte subtype classification*. Monocytes were identified by CD14 and CD45 co-expression. Subsets were classified based on CD14 and CD16 fluorescence intensity relative to control distributions. Classical monocytes were defined as CD14$^+$CD16$^-$, intermediate monocytes as CD14+CD16+, and non-classical monocytes as CD14$^{low}$CD16$^+$.

*Macrophage subtype classification*. Macrophages were classified based on HLA-DR and CD163 fluorescence intensity. M1-like macrophages were defined as HLA-DR$^{high}$CD163$^{low}$ and M2-like macrophages as HLA-DR$^{low}$CD163$^{high}$. The HLA-DR/CD163 ratio was calculated for each cell, and the threshold $>1.5$ separating M1 and M2 subsets (threshold $<1.5$) was determined from the distribution of this ratio across all macrophages.

*Proinflammatory immune phenotype score*. A composite proinflammatory immune phenotype score was defined as the combined proportion of Th17 cells, plasma cells, non-classical monocytes, and M1 macrophages among total classified immune cells for each chip.

**Scanning electron microscopy**

RA synovium-on-a-chip samples were cultured for 8 days, washed three times with PBS, and fixed with 2% glutaraldehyde in PBS (Electron Microscopy Sciences, 16010) for 1 hour. Fixed chips were manually sectioned into approximately 2-mm slices and dehydrated through a graded ethanol series (30%, 50%, 70%, 80%, and 90% for 10 minutes each, followed by three 20-minute washes in 100% ethanol). Samples were dried using a $CO_2$ critical point dryer (Samdri-PVT-3D, Tousimis), mounted on stubs, and sputter-coated with gold–palladium for 30 seconds (~12 nm coating). Images were acquired using a Gemini 300 FESEM with Gatan 3View (Zeiss).

**Single-cell suspension recovery from chips and synovial tissue**

For scRNA-seq analyses, single-cell suspensions were recovered from the synovium-on-a-chip by enzymatic digestion. Chips were washed with PBS and incubated with 100ug/ml Liberase TL (Cat: 05401020001) and 100ug/ml DNaseI (Cat: 10104159001) in RPMI at 37°C for 30 minutes with gentle pipetting to dissociate the fibrin hydrogel. The resulting cell suspensions were filtered through a 50 μm strainer, washed with cold PBS, centrifuged at 300 × g for 5 minutes at 4°C, and resuspended in PBS containing 0.04% BSA for downstream processing.

For scRNA-seq analyses, single-cell suspensions were recovered from synovial tissue specimens by enzymatic digestion. Cryopreserved tissue was thawed at 37°C, rinsed through a 40 µm strainer, and washed sequentially in RPMI with and without 10% FBS. Tissue was mechanically minced into a slurry and transferred into digestion medium containing Liberase TL (100 µg/mL) and DNase I (100 µg/mL) in serum-free RPMI. Samples were digested for 30 minutes at 37°C with continuous rotation. The resulting suspension was filtered through a 70 µm strainer, mashed with a syringe plunger, and washed with RPMI containing 10% FBS. Cells were pelleted at 1500 rpm for 4 minutes at 4°C, resuspended in cold RPMI with FBS, filtered through a 40 µm strainer, and counted using a Cellometer with AOPI viability staining.

**Gene set enrichment analysis**

Gene set enrichment analysis (GSEA) was performed using the fgsea (Fast Gene Set Enrichment Analysis) R package (version 1.36.2) with pre-ranked gene lists based on average log2 fold-change values from differential expression analyses. Hallmark gene sets (H collection) and Reactome canonical pathways (C2:CP:REACTOME) from msigdbr R package (version 25.1.1) were used as reference pathway databases. Gene sets with fewer than 15 or more than 500 genes were excluded. Enrichment significance was determined at an adjusted $P < 0.05$ (Benjamini–Hochberg correction). For cross-patient and cross-drug comparisons, normalized enrichment scores (NES) were compiled into heatmaps using the pheatmap R package. For pathway-level comparisons across FLS subpopulations (Fig. 5e), GSEA was performed independently for each cluster using pre-ranked gene lists derived from cluster-specific differential expression. Pathways were grouped into inflammation (TNFα/NF-κB signaling, inflammatory response), ECM (epithelial–mesenchymal transition, coagulation), and proliferation (E2F, MYC targets V1, G2M, mitotic spindle) categories for visualization.

**CellChat ligand–receptor interaction analysis**

Ligand–receptor interaction analysis was performed using CellChat (version 1.6) with the CellChatDB.human database. For each patient and treatment condition, a CellChat object was created from the Seurat object including all cell types on chip (eight FLS subpopulations, endothelial cells, lymphocytes, and macrophages). Overexpressed ligands and receptors were identified, communication probabilities were computed using the trimean method, and interactions with fewer than 5 cells in either the sender or receiver population were filtered. Signaling pathway-level communication probabilities were aggregated and compared across conditions. To identify signaling axes associated with treatment outcome, interaction probabilities for each sender–receiver pair and pathway were correlated with on-chip composite prediction scores across all six patient–drug conditions using Spearman rank correlation.

**FLS compositional and module score analysis**

For each patient and treatment condition, the proportion of cells in each of the eight FLS functional states was calculated relative to total FLS. Drug-induced compositional changes (Δ proportion) were defined as the difference between treated and untreated proportions for each state. To characterize FLS functional specialization, module scores were computed using the Seurat AddModuleScore function with the following gene sets: IFN-associated genes (ISG15, IFIT1, IFIT3, MX1, IFI6, IFITM3, OAS1), structural-remodeling-associated genes (SFRP1, SOX9, SCARA5, RIPOR2, ADH1B), ECM module genes (POSTN, ACTA2, THBS1, VCAN, COL1A1, COL3A1, FN1), and chemokine module genes (CCL2, CXCL12, CXCL1, CXCL8). For CD90-based FLS stratification, $CD90^+$ cells (normalized THY1 > 1.5) were stratified into CD90-low, CD90-mid, and CD90-high subpopulations by within-population tertiles of THY1 expression (cut-points 1.83 and 2.40). Double-positive cells (IFN score and structural-remodeling score both above the median) were classified as putative transition cells. To identify genes whose expression in $SFRP1^+$ FLS correlated with FN–CD44 signal strength, individual gene expression levels were correlated with the mean FN–CD44 interaction probability across patient–drug conditions using Pearson correlation, and the resulting ranked gene lists were used as input for GSEA.

**Composite chip treatment response score, clinical validation, and generation of predicted clinical response (**ΔDAS28-ESR)

Drug-induced changes (Δ values) were calculated by subtracting the untreated readout from the treated readout for each of four multimodal channels: immune cell migration distance (recruitment), IL-6 concentration (inflammation), CD45+ cell count (immune survival), and MMP secretion (ECM remodeling). For each patient and readout, treatment responses were expressed as treated-to-untreated ratios. Ratios for IL-6 secretion, $CD45^+$ cell count, and immune cell migration distance were inverted, whereas collagen IV ratios were retained. These direction-aligned ratios were z-score standardized across the three drug conditions separately for each readout within each patient, using the sample standard deviation. The four standardized values were averaged to obtain the composite on-chip response score. For clinical validation, composite prediction scores from five patients with documented prior TNF inhibitor and JAK inhibitor treatment were correlated with clinically observed ΔDAS28–ESR values (Pearson r = 0.884, p = 0.0348). A linear regression model (ΔDAS28 = 3.444 × score + 0.505) was derived from this correlation. Leave-one-out cross-validation (LOOCV) was performed by iteratively excluding each patient, fitting the model on the remaining three, and predicting the held-out patient's ΔDAS28. Root mean square error (RMSE) was calculated for each individual readout and the composite score to evaluate feature contribution. Applying established EULAR ΔDAS28 clinical treatment response criteria {van Gestel, 1996 #44}(good response: ΔDAS28 > 1.2; moderate response: </= 1.2 and > 0.6; no response: ΔDAS28 </= 0.6). Each patient's predicted response to each drug was classified as good, moderate, or no response. Composite chip treatment response and predicted clinical treatment response scores are depicted in Figure 5.

**Transcription factor activity inference**

Transcription factor (TF) activity in SFRP1⁺ FLS was inferred from log-normalized RNA expression using decoupleR (v2), the Homo sapiens DoRothEA regulons retrieved through OmnipathR, and the univariate linear model (ULM). Only high-confidence regulons (levels A–C) with at least five expressed targets were retained (minsize = 5). ULM-derived t-values were used as per-cell TF activity estimates. Per-cell activities were averaged within six patient–drug conditions (RA97, RA99, and RA105 treated with TNFi or JAKi), with the condition used as the unit of analysis (n = 6). TFs detected in fewer than five conditions were excluded. Mean TF activity was correlated with the CellChat-derived FN1–CD44 communication probability directed toward $SFRP1^+$ FLS in the corresponding condition using two-sided Spearman correlation. NFKB1 activity was negatively associated with FN1–CD44 signaling (R = −0.886, P = 0.019), whereas NR1H3/LXRα activity was positively associated (R = 0.943, P = 0.005). NR1H3 and NFKB1 activities were also inversely correlated (R = −0.943, P = 0.005).

**FN×CD44 imaging and quantification**

Chips were imaged by confocal microscopy using a 20X objective. Single optical planes were acquired as separate 16-bit TIFF images for DAPI, FN1, SFRP1, and CD44. 20 non-overlapping fields (>20 $SFRP1^+$ FLS) were collected per chip using identical acquisition settings without saturated pixels. Cells were segmented from the CD44 channel using Cellpose v3.1.1.3 (cyto3 model; expected diameter, 80 pixels) through the BIOP Cellpose wrapper in Fiji/ImageJ. Segmentation masks were converted to ROIs and manually curated by an operator blinded to treatment. Debris, out-of-focus objects, border-touching cells, and incorrectly segmented objects were excluded. Images were analyzed after rolling-ball background subtraction with a radius of 50 pixels. $SFRP1^+$ FLS were defined using a fixed whole-cell mean SFRP1 intensity threshold of ⩾100 a.u. A 10-pixel (3.2 μm) intracellular band adjoining the cell boundary was used to measure membrane-associated CD44, and a spatially matched 10-pixel extracellular ring was used to measure pericellular FN1. For each $SFRP1^+$ FLS, the FN×CD44 proximity index was calculated as: $\sqrt{FN \times CD44}$. Cell-level indices were averaged within each field. Field-level comparisons were performed using a two-sided Mann–Whitney U test and were considered exploratory because the fields were nested within individual chips.

### Supplementary Figures

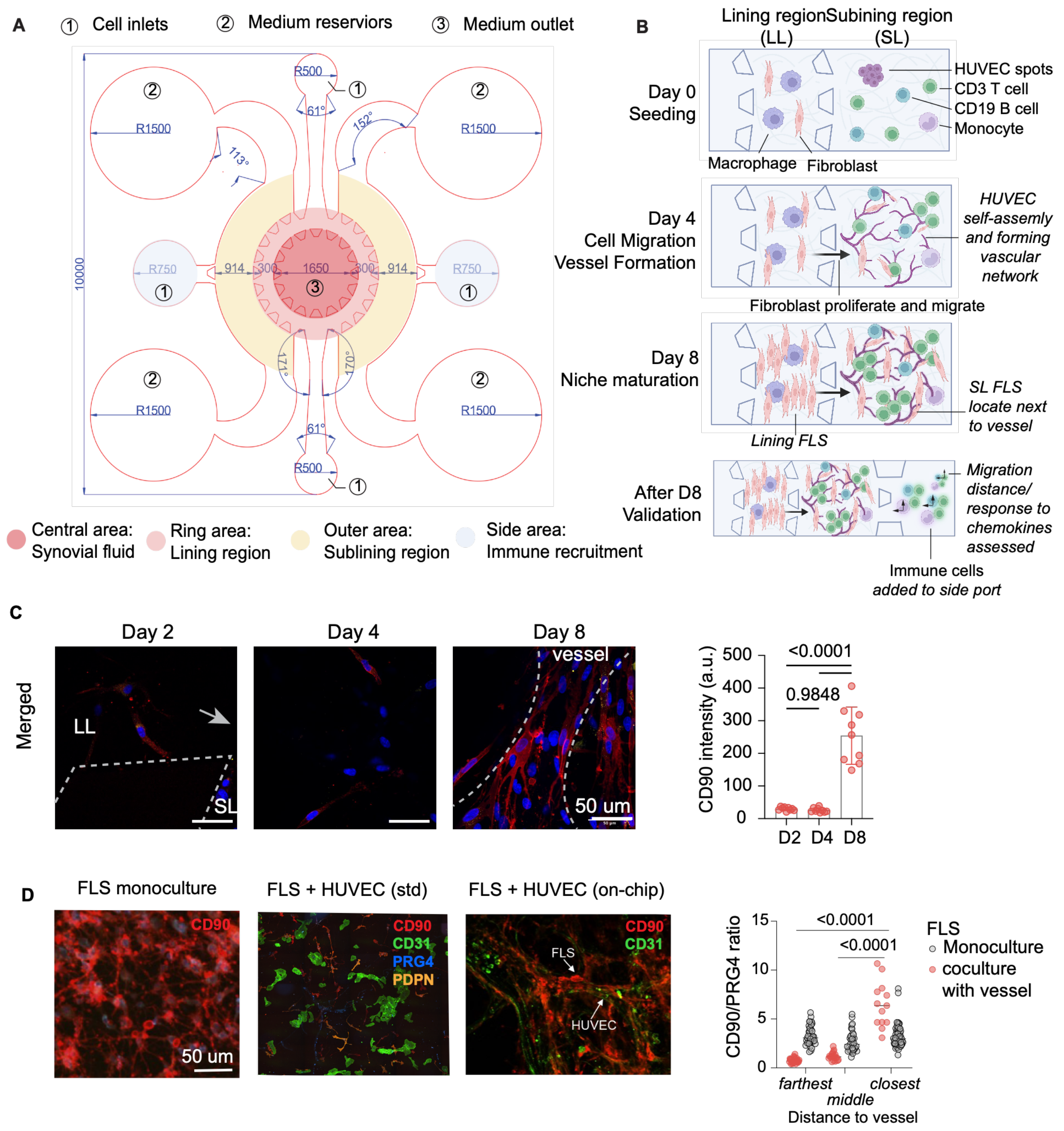


**Fig. S1. Chip assembly, spatial organization and temporal maturation of the RA synovium-on-a-chip platform**. (**A**) Microfluidic device layout illustrating cell inlets, medium reservoirs and outlet channels, with compartmentalized regions corresponding to synovial fluid (central), lining (ring), sublining (outer) and immune recruitment chambers (side). Device dimensions are indicated in micrometers. (**B**) Schematic of the cell loading and maturation timeline. At Day 0, fibroblast-like synoviocytes (FLS) and macrophages were seeded into the lining region, while HUVEC were introduced into the sublining compartment. By Day 4, HUVECs began to selfassemble into a vascular network and FLS began to proliferate and migrate toward the sublining region. By Day 8, the niche reached maturation with T cell proliferation, sublining FLS localizing adjacent to vessels, and lining FLS exhibiting hyperplasia. After Day 8, immune cell migration distance of cells from the same healthy donor in response to RA niche chemokine gradients was used as a functional validation readout. (**C**) Temporal monitoring of FLS spatial localization and differentiation on chip at Days 2, 4 and 8. Representative immunofluorescence images (merged channel, representative of n = 3 independent chips per time point) show progressive accumulation of $CD90^+$ fibroblasts from the lining layer (LL) toward the sublining (SL) and perivascular regions over time. Quantification of CD90 intensity demonstrates a significant increase by Day 8 compared to Day 2 and Day 4 (n = 3 chips; one-way ANOVA with multiple-comparison correction; $P < 0.0001$ for indicated comparisons; Day 2 vs Day 4, $P = 0.9848$). (**D**) Representative immunofluorescence images (n = 3 independent cultures per condition) of FLS monoculture on-chip, FLS coculture with HUVEC in a standard well plate, and FLS coculture with HUVEC on-chip. Monoculture and standard well plate cocultures show comparable CD90 signal intensity across the culture without perivascular enrichment. Quantification of the CD90/PRG4 ratio as a function of distance from the vessel region confirms no spatial gradient in monoculture, in contrast to the distance-dependent CD90 enrichment observed in vascular co-culture chips (right panel; n = 3 chips per condition, one-way ANOVA with multiple-comparison correction).

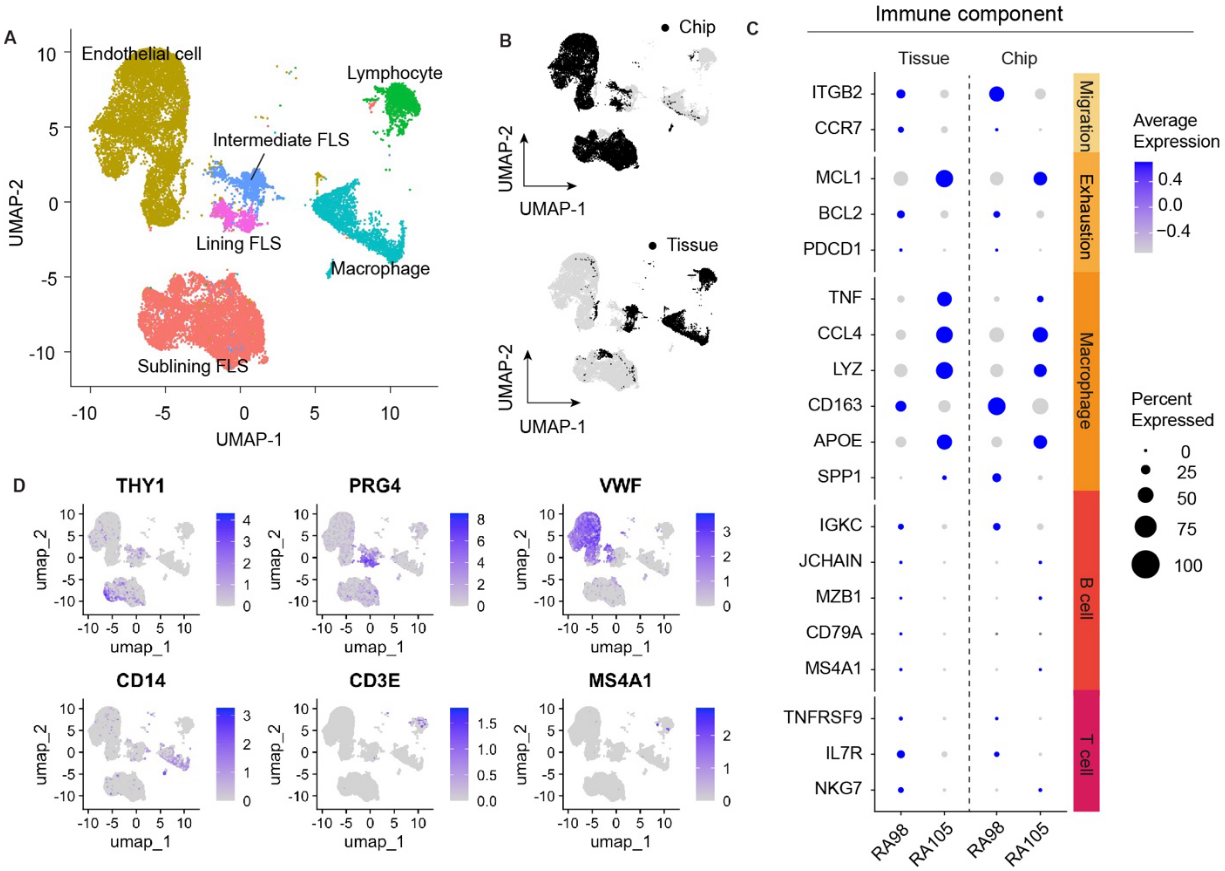


**Fig. S2. Transcriptional fidelity of the RA synovium-on-a-chip platform. (A)** UMAP visualization of scRNA-seq profiles from day-8 synovium-on-a-chip samples and matched synovial tissue from two patients: RA98 (high inflammatory endotype) and RA105 (low inflammatory endotype). Cells are colored by annotated population, including sublining FLS, lining FLS, intermediate FLS, endothelial cells, macrophages, and lymphocytes. **(B)** Distribution of chip-derived cells (top) and tissue-derived cells (bottom) within the UMAP embedding shown in panel A. Black points indicate cells from the specified source, and gray points indicate the remaining cells. **(C)** Dot plot showing selected immune-associated genes in matched tissue and chip samples from RA98 and RA105. Dot size indicates the percentage of cells expressing each gene, and color indicates scaled average expression. **(D)** UMAP feature plots showing the expression of selected cell-type markers: THY1, PRG4, VWF, CD14, CD3E, and MS4A1. Color intensity indicates normalized gene expression.

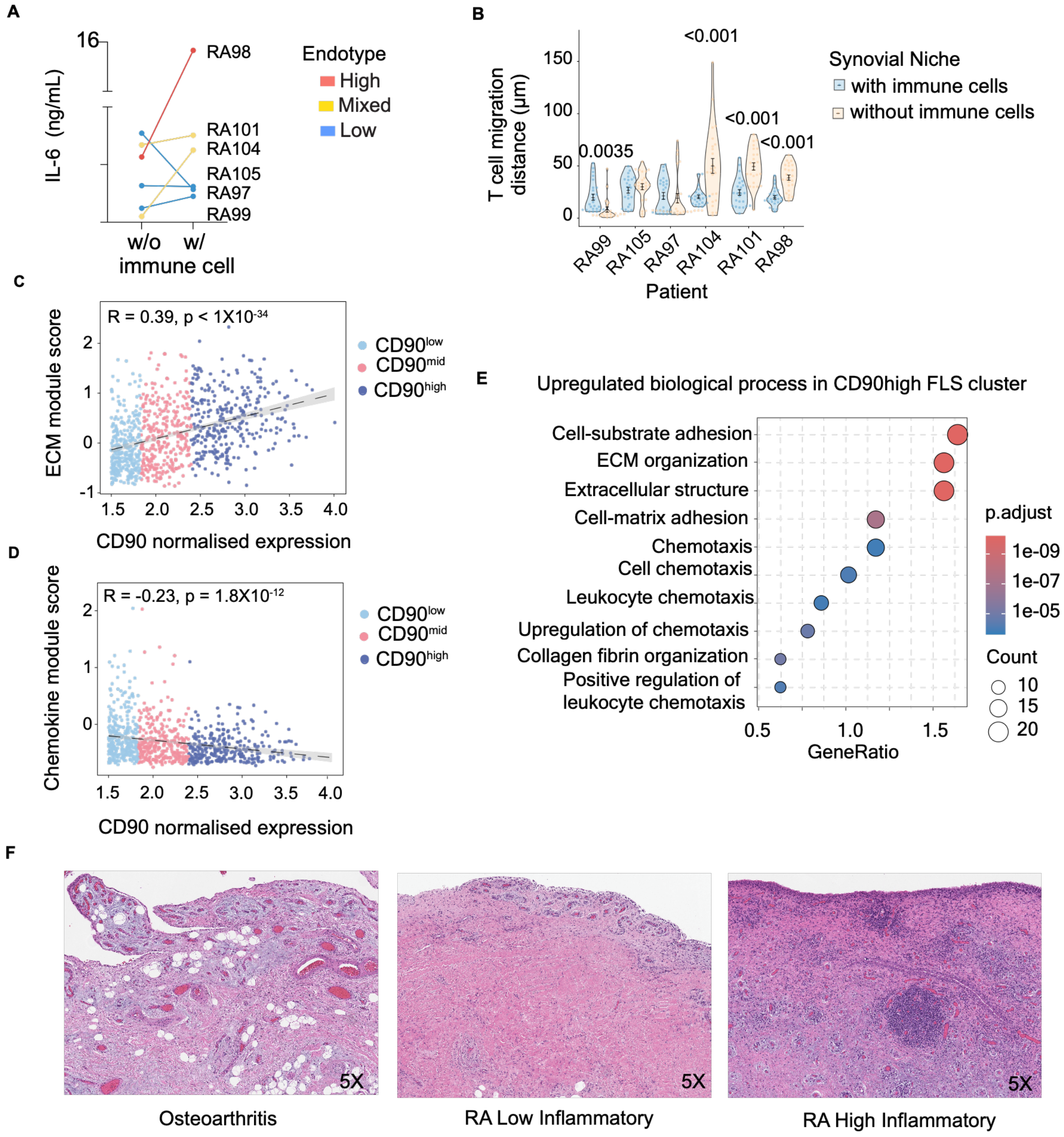


**Fig. S3. Stromal-immune crosstalk. (A)** IL-6 secretion in synovium-on-a-chips created with or without immune cells (T cells, B cells, or monocytes) across individual patients, showing patient-specific changes in inflammatory cytokine production following immune-cell incorporation. **(B)** Distribution of T cell migration distances in synovium-on-a-chips created with or without immune cells (T cells, B cells, or monocytes) across individual patients (RA99, RA105, RA97, RA104, RA101, and RA98). Immune-cell incorporation altered T cell migration in a patient-dependent manner (Wilcoxon rank-sum test; $P = 0.0035$ for RA99 and $P < 0.001$ for RA104, RA101, and RA98). **(C)** Correlation between normalized CD90 expression and ECM module score across individual FLS, colored by CD90$^{low}$, CD90$^{mid}$, and CD90$^{high}$ subpopulations. ECM module scores increased with CD90 expression

(Pearson R = 0.39, $P < 1 \times 10^{-34}$). **(D)** Correlation between normalized CD90 expression and chemokine module score across individual FLS, showing an inverse relationship between chemokine programs and CD90 expression (Pearson R = −0.23, $P = 1.8 \times 10^{-12}$). **(E)** Gene Ontology biological process analysis of genes upregulated in the $CD90^{high}$ FLS cluster, showing enrichment of ECM organization and cell–matrix adhesion programs together with chemotaxis- and leukocyte recruitment-associated processes. Dot size represents the number of associated genes, and color indicates the adjusted P value. **(F)** Representative H&E-stained synovial tissue sections from osteoarthritis (disease comparison/control), the RA low inflammatory endotype showing FLS abundance with loss of adipose, and the high-inflammatory endotype, showing dense inflammatory-cell accumulation.

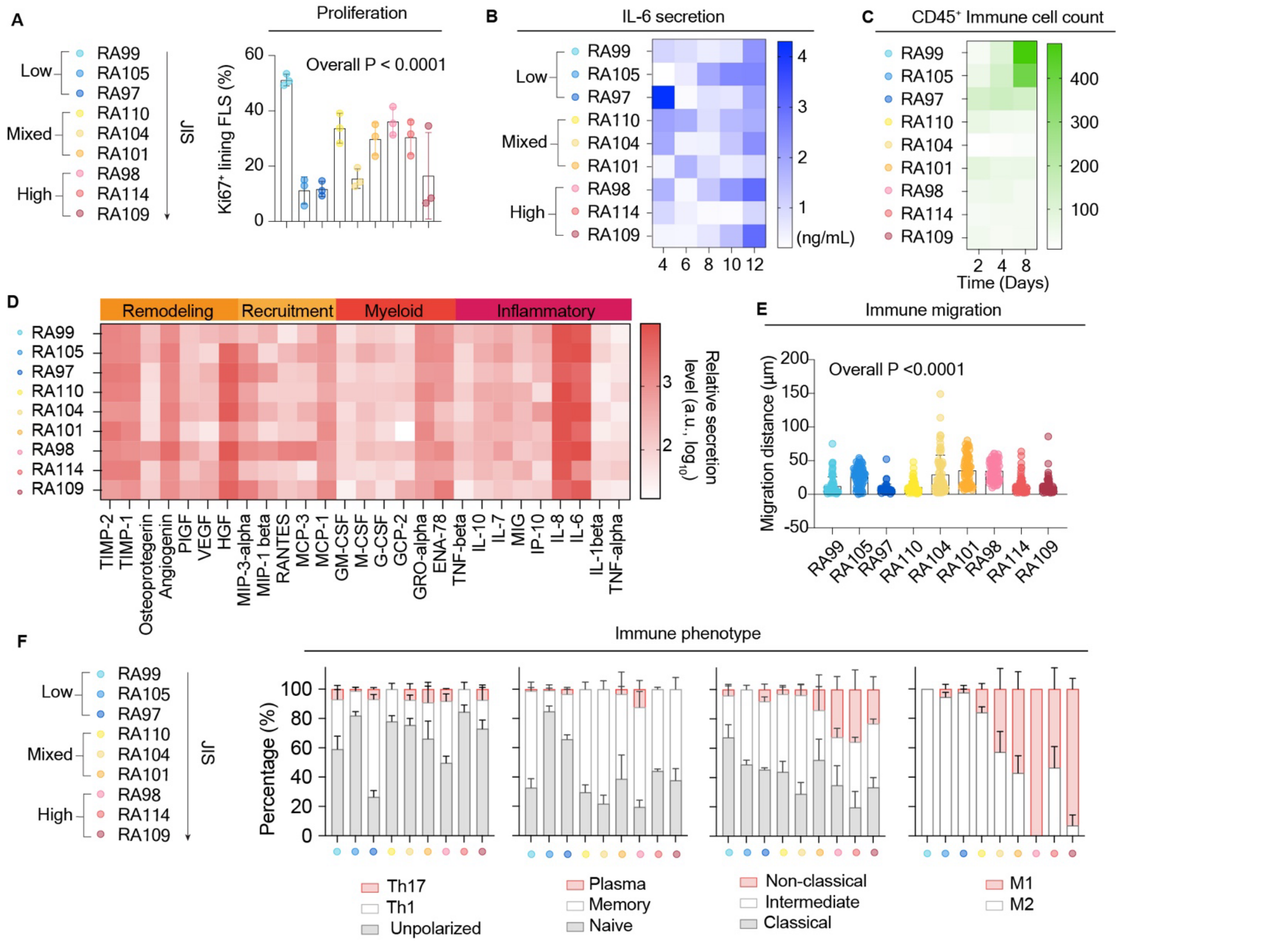


**Fig. S4. Patient-level inflammatory dynamics, FLS subset kinetics and clinical correlations in RA synovium-on-a-chip. (A)** Percentage of Ki67⁺ lining FLS across nine patients grouped by synovial inflammatory endotype: low (RA99, RA105 and RA97), mixed (RA110, RA104 and RA101), and high (RA98, RA114 and RA109). Bars and error bars indicate mean ± SD. Each dot represents one chip. Differences among patients were assessed using ordinary one-way ANOVA ($P < 0.0001$). **(B)** Heatmap showing IL-6 concentrations in chip culture supernatants on Days 4, 6, 8, 10 and 12 across individual patients. Color indicates IL-6 concentration (ng/mL). **(C)** Heatmap showing on-chip CD45⁺ immune cell counts on Days 2, 4 and 8 across individual patients. **(D)** Heatmap of secreted factors associated with tissue remodeling, immune recruitment, myeloid function and inflammation. Color indicates $\log_{10}$-transformed relative secretion levels in arbitrary units. Measurements were obtained on Day 12. **(E)** Immune cell migration distances across nine patients. Each dot represents one tracked cell. Cells were analyzed >75 cells from 3 technical repeats per patient; bars and error bars indicate mean ± SD. Differences among patients were assessed using ordinary one-way ANOVA ($P < 0.0001$). **(F)** Immune cell phenotypic composition across individual patients grouped by inflammatory endotype and ordered by increasing JIS. Stacked bars show the proportions of T-cell subsets (Th17, Th1 and unpolarized), B-cell subsets (plasma, memory and naive), monocyte subsets (non-classical, intermediate and classical), and macrophage subsets (M1 and M2) within their respective parent populations. Data are presented as mean ± SEM across 3 chips per patient. For (B–D), each heatmap value represents mean across 3 chips per patient per condition.

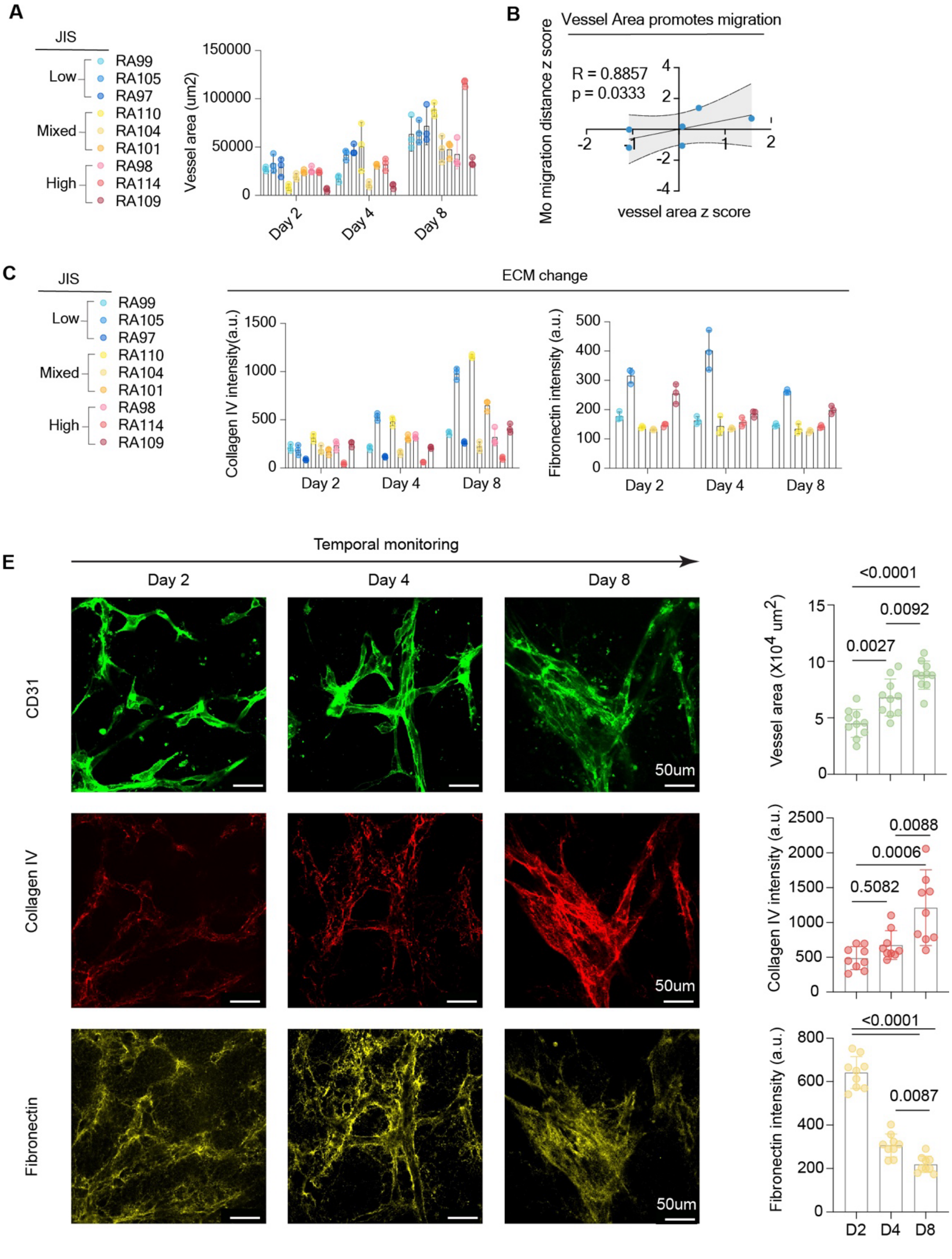


**Fig S5. Patient-level vascular and ECM dynamics and their clinical correlations in the RA synovium-on-a-chip.** (**A**) Vessel area quantified for each individual patient at Days 2, 4 and 8, stratified by inflammatory endotype (low, mixed and high). (**B**) Vessel area (z-score normalized) positively correlated with monocyte migration distance (z-score normalized) across patients (R = 0.8857, p = 0.0333; Pearson correlation, two-tailed). Each dot represents an individual patient. (**C**) Collagen IV intensity quantified for each individual patient at Days 2, 4 and

8, stratified by inflammatory endotype. (**D**) Temporal monitoring of ECM and vascular remodeling at Days 2, 4 and 8. Representative immunofluorescence images show progressive expansion of $CD31^+$ vascular networks (top, green), dynamic changes in collagen IV deposition (middle, red) and fibronectin remodeling (bottom, yellow). Quantification revealed significant increases in vessel area over time (one-way ANOVA with multiple-comparison correction; Day 2 vs Day 4, $p = 0.0027$; Day 4 vs Day 8, $p = 0.0092$; overall $p < 0.0001$). Collagen IV intensity increased significantly between Day 2 and Day 8 ($p = 0.0006$) and Day 4 and Day 8 ($p = 0.0088$), with no significant change between Day 2 and Day 4 ($p = 0.5082$; overall $p < 0.0001$). Fibronectin intensity decreased significantly over time (Day 4 vs Day 8, $p = 0.0087$; overall $p < 0.0001$).

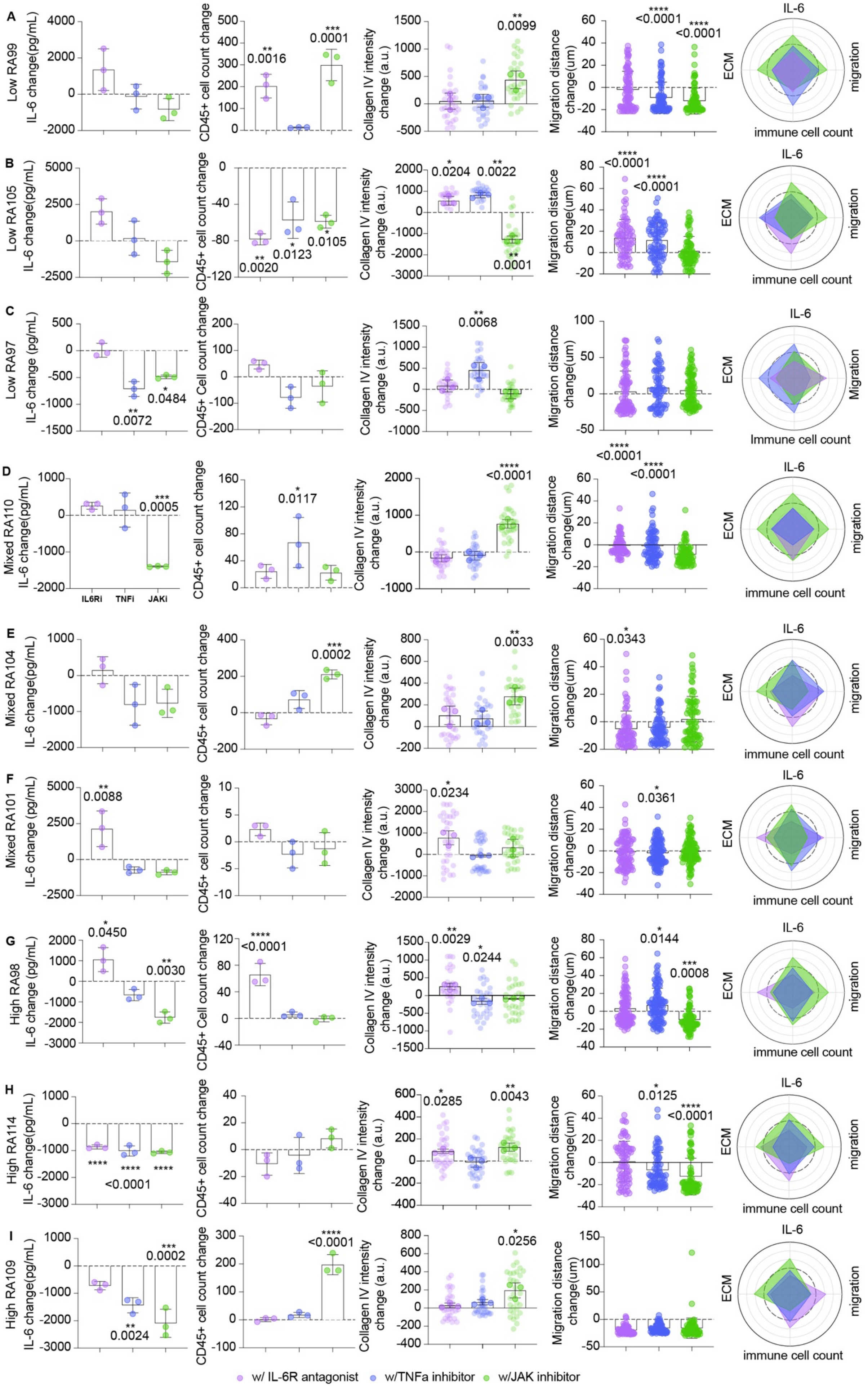

A
Low RA99
IL-6 change(pg/mL)
CD45+ cell count change
0.0016
0.0001
Collagen IV intensity change (a.u.)
0.0099
Migration distance change(um)
<0.0001
<0.0001
IL-6
ECM
migration
immune cell count
B
Low RA105
IL-6 change(pg/mL)
CD45+ cell count change
0.0020
0.0123
0.0105
Collagen IV intensity change (a.u.)
0.0204
0.0022
0.0001
Migration distance change(um)
<0.0001
<0.0001
C
Low RA97
IL-6 change (pg/mL)
0.0072
0.0484
CD45+ Cell count change
Collagen IV intensity change (a.u.)
0.0068
Migration distance change(um)
Migration
Immune cell count
D
Mixed RA110
IL-6 change(pg/mL)
0.0005
IL6Ri
TNFi
JAKi
CD45+ cell count change
0.0117
Collagen IV intensity change (a.u.)
<0.0001
Migration distance change(um)
<0.0001
<0.0001
E
Mixed RA104
IL-6 change(pg/mL)
CD45+ cell count change
0.0002
Collagen IV intensity change (a.u.)
0.0033
Migration distance change(um)
0.0343
F
Mixed RA101
IL-6 change (pg/mL)
0.0088
CD45+ cell count change
Collagen IV intensity change (a.u.)
0.0234
Migration distance change(um)
0.0361
G
High RA98
IL-6 change (pg/mL)
0.0450
0.0030
CD45+ Cell count change
<0.0001
Collagen IV intensity change (a.u.)
0.0029
0.0244
Migration distance change(um)
0.0144
0.0008
H
High RA114
IL-6 change(pg/mL)
<0.0001
CD45+ cell count change
Collagen IV intensity change (a.u.)
0.0285
0.0043
Migration distance change(um)
0.0125
<0.0001
I
High RA109
IL-6 change(pg/mL)
0.0002
0.0024
CD45+ cell count change
<0.0001
Collagen IV intensity change (a.u.)
0.0256
Migration distance change(um)
w/ IL-6R antagonist
w/TNFa inhibitor
w/JAK inhibitor

**Fig S6. Patient-specific drug responses and multivariate treatment outcome prediction on RA synovium-on-a-chip.** (**A–I**) Changes in inflammatory and tissue remodeling features following treatment with tocilizumab (IL-6 receptor antagonist, IL-6Ra), infliximab (TNF inhibitor, TNFi), or tofacitinib (JAK inhibitor, JAKi) in synovium-on-a-chips derived from individual RA patients. Panels show chips derived from patients RA99 (A), RA105 (B), and RA97 (C), classified as low inflammatory endotype; RA110 (D), RA104 (E), and RA101 (F), classified as mixed inflammatory endotype; and RA98 (G), RA114 (H), and RA109 (I), classified as high inflammatory endotype. Changes were calculated as treated-untreated, with zero indicating no change. Measurements were obtained at Day 12. Radar plots visualize the four-feature response profiles for each patient and treatment. Purple, blue, and green indicate IL-6Ra, TNFi, and JAKi treatment, respectively. Three independently cultured chips were analyzed per patient per condition. For IL-6 secretion, each point represents one chip. For $CD45^+$ cell counts, each point represents one chip. For collagen IV intensity, lighter points represent individual imaging fields (10 fields per chip); field measurements were averaged within each chip, and the three chip-level means were used for statistical analysis. Migration distances were measured in >75 cells across three technical replicates; each point represents one cell. Bars and error bars indicate mean ± SD. All statistical analyses were compared between each treatment and the untreated control using ordinary one-way ANOVA followed by Dunnett's multiple-comparisons test, with three chips per group. Adjusted P values are shown for comparisons with the untreated control.

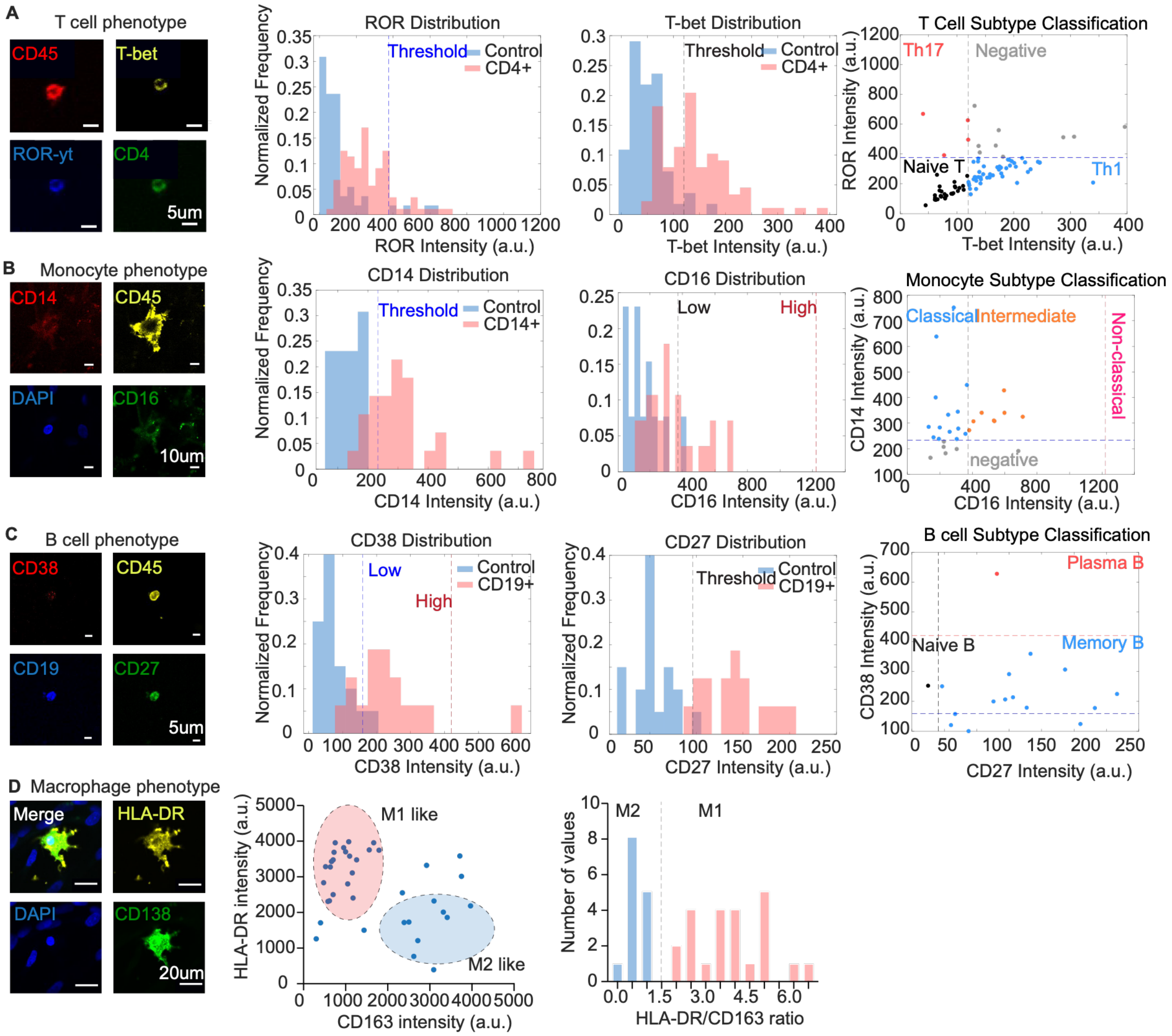


**Fig. S7. Immune cell phenotyping and inflammatory dynamics in RA synovium-on-a-chip. (A)** T cell subtype classification on chip. Representative immunofluorescence images showing CD45, T-bet, ROR-γt and CD4 staining (left). Normalized frequency distributions of ROR-γt and T-bet intensity in $CD4^+$ cells versus unlabeled controls, with dashed lines indicating classification thresholds (middle). Scatter plot of ROR-γt versus T-bet intensity per cell, stratified into Th17 (ROR-$γt^+$), Th1 (T-$bet^+$) and naive T cell subsets (right). **(B)** Monocyte subtype classification on chip. Representative immunofluorescence images showing CD14, CD45, DAPI and CD16 staining (left). Normalized frequency distributions of CD14 and CD16 intensity in $CD14^+$ cells versus controls, with dashed lines indicating thresholds (middle). Scatter plot of CD14 versus CD16 intensity per cell, stratified into classical ($CD14^+CD16^–$), intermediate ($CD14^+CD16^+$) and non-classical ($CD14lowCD16^+$) monocyte subsets (right). **(C)** B cell subtype classification on chip. Representative immunofluorescence images showing CD38, CD45, CD19 and CD27 staining (left). Normalized frequency distributions of CD38 and CD27 intensity in $CD19^+$ cells versus controls, with dashed lines indicating thresholds (middle). Scatter plot of CD38 versus CD27 intensity per cell, stratified into plasma B cells (CD38high), memory B cells ($CD27^+$CD38low) and naive B cells ($CD27^-$CD38low) subsets (right). (**D**) Macrophage subtype classification on chip. Representative immunofluorescence images showing merged, HLA-DR, DAPI and CD163 staining (left). Scatter plot of HLA-DR versus CD163 intensity per cell, identifying M1-like (HLA-DRhighCD163low) and M2-like (HLA-DRlowCD163high) populations (middle). Distribution of HLA-DR/CD163 ratio across macrophages, with dashed line (1.5) indicating the threshold separating M1 and M2 subsets (right).

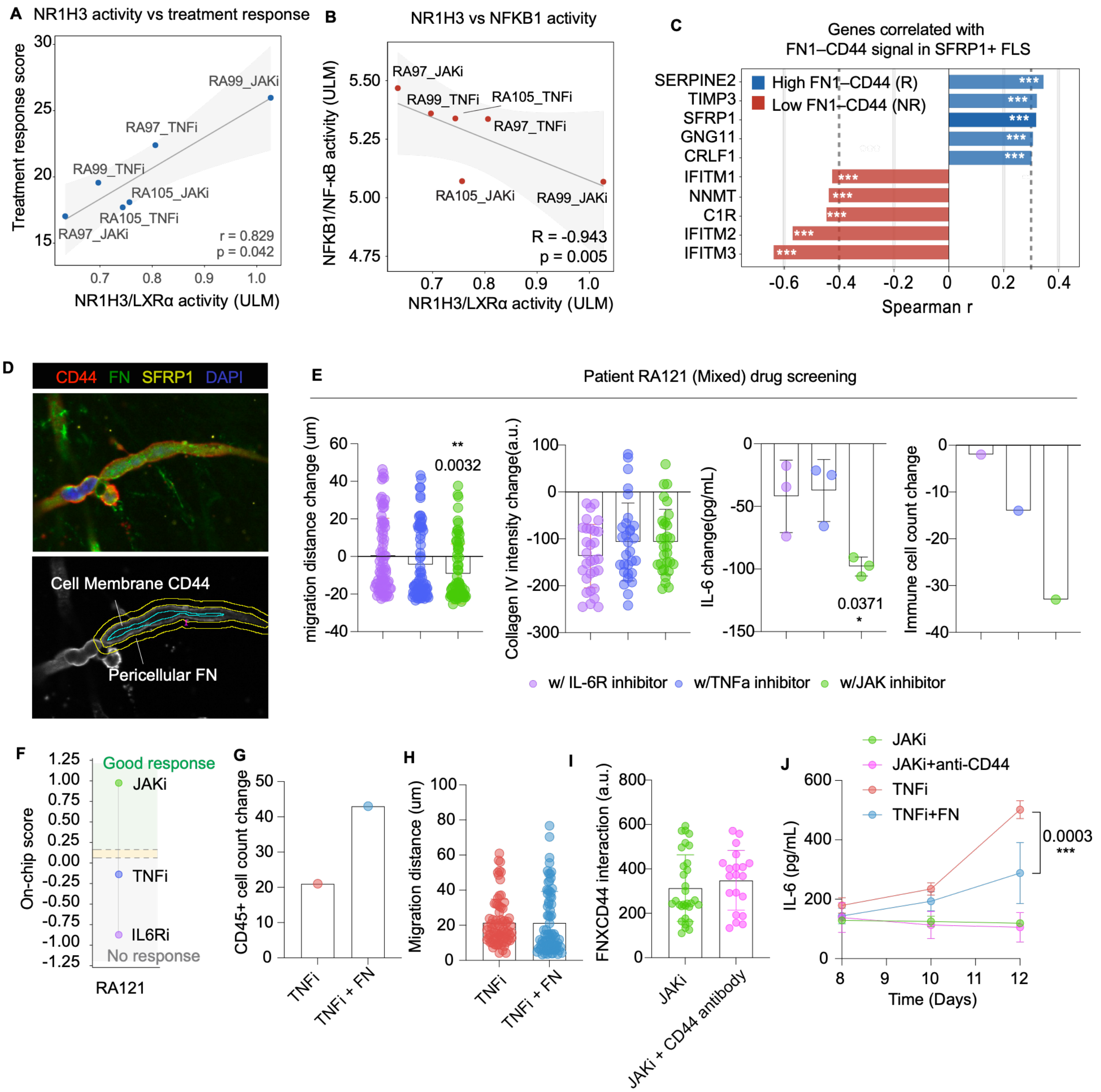


**Fig. S8. NR1H3–NF-κB activity and functional evaluation of FN–CD44 niche modulation. (A)** Correlation between inferred NR1H3/LXRα activity in SFRP1$^{+}$ FLS and the composite on-chip treatment response score across six patient–treatment conditions. Higher NR1H3 activity was associated with better treatment response (r = 0.829, P = 0.042). Each dot represents one patient–treatment condition; **(B)** Inverse relationship between NR1H3/LXRα and NFKB1/NF-κB activities across the same conditions, consistent with opposing LXRα-high/NF-κB-low and LXRα-low/NF-κB-high transcriptional states (R = −0.943, P = 0.005). **(C)** Spearman correlation analysis between FN1–CD44 signaling strength and gene expression in SFRP1$^{+}$ FLS across treated conditions. *SERPINE2, TIMP3, SFRP1, GNG11*, and *CRLF1* were positively associated with the high-FN1–CD44 responder state, whereas *IFITM1, NNMT, C1R, IFITM2*, and *IFITM3* were negatively associated with FN1–CD44 signaling and characterized the low-FN1–CD44 nonresponder state. Bars represent Spearman correlation coefficients. $P < 0.001$. **(D)** Representative immunofluorescence image showing the spatial association of CD44, fibronectin (FN), and SFRP1 within the synovium-on-a-chip (top), and illustration of the image-analysis strategy used to quantify FN–CD44 interactions (bottom). A five-pixel-wide (1.6 μm) region surrounding the $CD44^+$ cell membrane was defined as the pericellular FN interaction zone, and the pixel-wise product of CD44 and FN fluorescence within this region was used to calculate the FN×CD44 interaction score. **(E)** Multimodal drug screening in chips derived from patient RA121, classified as having a mixed inflammatory endotype. Treatment-induced changes in immune cell migration distance, collagen IV intensity, IL-6 secretion, and $CD45^+$ immune cell count are shown following IL-6 receptor antagonist (IL-6Ra), TNF inhibition (TNFi), or JAK inhibition (JAKi) relative to untreated controls. P values are indicated for significant comparisons. **(F)** Integrated on-chip treatment response scores for RA121, predicted a good response to JAKi, no response to TNFi, and no response to IL-6Ra. Background shading indicates the predicted response categories. **(G)** Change in $CD45^+$ immune cell count following treatment with TNFi alone or TNFi combined with exogenous fibronectin (TNFi + FN). **(H)** Distribution of immune cell migration distances following TNFi or TNFi + FN treatment. Each dot represents one quantified field. **(I)** FN×CD44 interaction signals following JAKi treatment alone or in combination with an anti-CD44 blocking antibody, showing no appreciable reduction in the interaction signal following CD44 blockade. Each dot represents one quantified field. **(J)** Temporal IL-6 secretion from Days 8 to 12 following JAKi, JAKi + anti-CD44, TNFi, or TNFi + FN treatment. Fibronectin supplementation significantly attenuated the increase in IL-6 secretion observed with TNFi alone, whereas anti-CD44 treatment had minimal additional effect on JAKi response ($P = 0.0003$ at Day 12). Data are shown as mean with error bars as indicated.

**Supplementary Table 1. Clinical characteristics of RA study participants.** Summary of patient histopathological inflammatory endotype classification (low, mixed, or high); joint inflammation score (JIS); tender joint count (TJC), swollen joint count (SJC) and disease activity scores (DAS28-CRP and DAS28-ESR) at the time of sample acquisition (DOS: date of surgery); rheumatoid factor (RF) and anti-citrullinated (CCP) antibody serostatus (positive or negative); disease modifying antirheumatic drug treatment history; and actual treatment response (EULAR response categorization provided when sufficient data was available). Timeframe in parenthesis denotes duration of therapy at which time the response was noted (mo: month, yr: year). Inflammatory endotype was determined from the joint inflammation score (JIS), as described in the Methods. A total of nine RA patients (RA97–RA109) were included for platform characterization, endotype analysis, and on-chip drug response profiling; a subset of five patients was used for correlation with real-world treatment response, as these patients had real world treatment response data available for therapy categories tested on-chip; three low-endotype patients (RA97, RA99, RA105) with divergent drug response profiles were selected for single-cell RNA sequencing. An additional patient (RA121) was included for functional examination of treatment resistant niche modulation.

| Patient | Inflammation Endotype | Joint inflammation score (JIS) | DOS TJC | DOS SJC | DOS DAS28-CRP | DOS DAS28-ESR | RF | CCP antibody | Treatment History | | Treatment Response (EULAR GR, MR, NR where available) |
|---|---|---|---|---|---|---|---|---|---|---|---|
| | | | | | | | | | Medication | Drug Class | |
| RA97 | Low | 1.6 | 12 | 3 | 2.86 | 3.59 | - | + | adalimumab | TNFi (bDMARD) | GR to TNFi (2, 6, 9 mo) |
| RA99 | Low | 0.8 | 3 | 1 | 3.43 | 4.37 | + | + | Mtx | cDMARD | NR (1-8 mo) |
| | | | | | | | | | Mtx + adalimumab | cDMARD + TNFi (bDMARD) | NR (5, 7 mo) |
| | | | | | | | | | Mtx + etanercept | cDMARD + TNFi (bDMARD) | MR (4, 10 mo) |
| | | | | | | | | | Mtx + rituximab | cDMARD + CD20 B cell depletion (bDMARD) | MR (4 mo) |
| RA105 | Low | 1.2 | 1 | 1 | 2.23 | 3.17 | - | - | Mtx + golimumab | cDMARD + TNFi (bDMARD) | NR (1, 3, 6, 12, 18 mo) |
| RA101 | Mixed | 6.4 | 5 | 3 | 2.48 | 3.76 | + | + | Mtx | cDMARD | NR (6-18 mo) |
| | | | | | | | | | rituximab | CD20 B cell deplation (bDMARD) | MR (4 mo) |
| RA104 | Mixed | 5.1 | 2 | 0 | 2.12 | 2.77 | + | - | Mtx | cDMARD | MR (3 mo) |
| RA110 | Mixed | 4.7 | 2 | 2 | | | + | + | abatacept | CTLA-4-IgG1 Fc (bDMARD) | No change in SJC and TJC (12+ mo) |
| RA98 | High | 7.4 | 1 | 4 | 3.34 | 5.06 | + | + | adalimumab | TNFi (bDMARD) | NR to TNFi (1, 2 mo) |
| RA109 | High | 9.8 | 2 | 2 | 4.37 | 4.89 | + | + | Mtx | cDMARD | GR (2, 5, 9 mo), MR (4, 13, 20 mo), NR (16 mo) |
| RA114 | High | 7.9 | 9 | 8 | 4.12 | 4.99 | + | + | Mtx + adalimumab | cDMARD + TNFi (bDMARD) | NR (2, 3 mo) |
| | | | | | | | | | Mtx + tofacitinib | cDMARD + JAKi (tsDMARD) | MR (12 mo) |
| | | | | | | | | | tocilizumab | IL6Ri (bDMARD) | Stopped due to infection (2 mo) |
| | | | | | | | | | golimumab | TNFi (bDMARD) | Improved SJC & TJC (7 mo), worsened SJC & TJC than pre-2nd TNFi (9 mo) |
| | | | | | | | | | infliximab | TNFi (bDMARD) | Improved SJC & TJC (3 mo) |
| RA121 | Mixed | 4.8 | 4 | 9 | 4.02 | 4.3 | + | + | abatacept | CTLA-4-IgG1 Fc (bDMARD) | No response (3 mo) |

**Supplementary Table 2.** Antibodies used for immunofluorescence staining of synovium-on-a-chip.

| Antibody | Fluorescence | Dilution | Host | Cat# | Manufacturer |
|---|---|---|---|---|---|
| DAPI | Blue | 1 µg/mL | - | D1306 | Thermo Fisher Scientific |
| CD31 | FITC | 1:50 | Mouse | 303104 | Biolegend |
| Podoplanin (PDPN) | PE | 1:20 | Rat | NBP203952JF549 | Novus Biologicals |
| CD3 | APC | 1:50 | Mouse | 317318 | Biolegend |
| CD4 | PE | 1:50 | Mouse | 317410 | Biolegend |
| CD8 | FITC | 1:50 | Mouse | 344704 | Biolegend |
| CD3 | PE | 1:50 | Mouse | 317308 | Biolegend |
| RORγt | Alexa Fluor® 647 | 1:50 | Mouse | 563620 | BD Pharmingen |
| T-bet | PE | 1:50 | Mouse | 644809 | Biolegend |
| CD19 | FITC | 1:50 | Mouse | 302206 | Biolegend |
| CD27 | FITC | 1:50 | Mouse | 356404 | Biolegend |
| CD38 | Brilliant Viole 421 | 1:50 | Mouse | 356618 | Biolegend |
| Ki-67 | FITC | 1:50 | Mouse | 350508 | Biolegend |
| Fibronectin | PE | 1:50 | Mouse | IC1918P | R&D Systems |
| Collagen IV | Alexa Fluo 647 | 1:50 | Mouse | 51-9871-80 | Biolegend |
| CD163 | Alexa Fluor 647 | 1:50 | Mouse | 333622 | Biolegend |
| HLA-DR | Alexa Fluor 488 | 1:50 | Mouse | 307640 | Biolegend |
| CD90 | APC | 1:50 | Mouse | 328113 | Biolegend |
| FAP | PE | 1:50 | Mouse | FAB3715P-025 | R&D Systems |
| CD55 | Alexa Fluor® 546 | 1:50 | Mouse | sc-51733 AF546 | Santa Cruz |
| PRG4 | - | 1:50 | Rabbit | ZRB1643-25UL | Sigma-Aldrich |
